# Tumor microenvironmental determinants of high-risk DCIS progression

**DOI:** 10.1101/2023.12.01.569676

**Authors:** Alexa Glencer, Kirithiga Ramalingam, Nicole Schindler, Hidetoshi Mori, Prachi Ghule, Kyra Lee, Daniela Nachmanson, Adam Officer, Olivier Harismendy, Janet Stein, Gary Stein, Mark Evans, Donald Weaver, Christina Yau, Gillian L Hirst, Michael J Campbell, Laura J Esserman, Alexander D. Borowsky

## Abstract

Ductal carcinoma *in situ* (DCIS) constitutes an array of morphologically recognized intraductal neoplasms in the mammary ductal tree defined by an increased risk for subsequent invasive carcinomas at or near the site of biopsy detection. However, only 15-45% of untreated DCIS cases progress to invasive cancer, so understanding mechanisms that prevent progression is key to avoid overtreatment and provides a basis for alternative therapies and prevention. This study was designed to characterize the tumor microenvironment and molecular profile of high-risk DCIS that grew to a large size but remained as DCIS. All patients had DCIS lesions >5cm in size with at least one additional high-risk feature: young age (<45 years), high nuclear grade, hormone receptor negativity, HER2 positivity, the presence of comedonecrosis, or a palpable mass. The tumor immune microenvironment was characterized using multiplex immunofluorescence to identify immune cells and their spatial relationships within the ducts and stroma. Gene copy number analysis and whole exome DNA sequencing identified the mutational burden and driver mutations, and quantitative whole-transcriptome/gene expression analyses were performed. There was no association between the percent of the DCIS genome characterized by copy number variants (CNAs) and recurrence events (DCIS or invasive). Mutations, especially missense mutations, in the breast cancer driver genes *PIK3CA* and *TP53* were common in this high-risk DCIS cohort (47% of evaluated lesions). Tumor infiltrating lymphocyte (TIL) density was higher in DCIS lesions with TP53 mutations (p=0.0079) compared to wildtype lesions, but not in lesions with *PIK3CA* mutations (p=0.44). Immune infiltrates were negatively associated with hormone receptor status and positively associated with HER2 expression. High levels of CD3+CD8-T cells were associated with good outcomes with respect to any subsequent recurrence (DCIS or invasive cancer), whereas high levels of CD3+Foxp3+ Treg cells were associated with poor outcomes. Spatial proximity analyses of immune cells and tumor cells demonstrated that close proximity of T cells with tumor cells was associated with good outcomes with respect to any recurrence as well as invasive recurrences. Interestingly, we found that myoepithelial continuity (distance between myoepithelial cells surrounding the involved ducts) was significantly lower in DCIS lesions compared to normal tissue (p=0.0002) or to atypical ductal hyperplasia (p=0.011). Gene set enrichment analysis identified several immune pathways associated with low myoepithelial continuity and a low myoepithelial continuity score was associated with better outcomes, suggesting that gaps in the myoepithelial layer may allow access/interactions between immune infiltrates and tumor cells. Our study demonstrates the immune microenvironment of DCIS, in particular the spatial proximity of tumor cells and T cells, and myoepithelial continuity are important determinants for progression of disease.

## INTRODUCTION

Ductal carcinoma in situ (DCIS) is a subset of intraductal proliferative breast disease defined by an increased risk of subsequent invasive carcinoma at or near the site of the biopsy/excision^1^. Although heterogeneous, DCIS is characterized by clonal proliferation replacing the normal luminal epithelium and with architectural and often cytologic atypia confined within the basement membrane and specialized periductal stroma^2^. The routine use of mammography for breast cancer screening over the past thirty years has increased incidence of these lesions from 3% to greater than 25% of all breast cancers detected^3^. Standard treatments for DCIS are complete surgical resection, radiation and endocrine therapy for hormone receptor (HR)-positive disease, based upon the assumption that any/all DCIS lesions have potential for progression to invasive carcinomas. However, DCIS represents a spectrum of biologic risk^4,5^ and a uniform approach to the management of DCIS has not led to an overall reduction in invasive breast cancer. This suggests that both overtreatment of indolent lesions and inadequate treatment of more aggressive lesions are contributing factors^6,7^.

Clinical characteristics such as young age and tumor characteristics such as large size, high-grade, HR-negative status, HER2-positive status, the presence of comedonecrosis, and palpability have each been shown to predict increased risk of DCIS progression to invasive breast cancer^8–10^. Yet despite identification of these risk factors, much remains unknown about the biology of high-risk DCIS.

Some studies have suggested that lesions with documented/synchronous progression have similar genomic profiles to invasive disease^11,12^, but others have demonstrated that genomic alterations occur with the progression of DCIS to invasive cancer^13,14^. In the DCIS tumor immune microenvironment, specific immune cell infiltrate patterns have been associated with high-risk clinical features^15^. The proximity of these populations to one another and localization within ducts versus adjacent and peripheral stromal compartments has potential utility in predicting progression^16,17^. The density of tumor infiltrating lymphocytes in the immune microenvironment has also been associated with certain gene expression profiles, which has potential implications for the development of targeted therapeutics^18^.

In this study, we assembled a cohort of high-risk DCIS cases, defined by large size (>5cm) and at least one additional clinicopathologic attribute associated with high risk of progression to invasive disease in order to enrich for incidence of invasive progression but also to better understand the biology of lesions which grow very large yet remain as DCIS. We characterized the genomic and tumor immune microenvironment characteristics of this “DEFENSE” cohort (DCIS: Elaboration of Factors from Enlarged lesions that Nevertheless remain Stage 0 Entities) and investigated associations with DCIS recurrence or development of invasive breast cancer. All high-risk DCIS in this cohort were treated with standard of care excision, radiation, and/or hormonal therapy, yielding rates of recurrence or progression to invasive breast cancer in the context of treatment. Without accurate predictors of disease progression, many patients with DCIS undergo surgical resection with no survival benefit, while others with biologically aggressive disease are undertreated and would benefit from targeted molecular and immune therapy. Understanding the underlying genomic profile of high-risk DCIS and the interaction between these lesions and their immune microenvironment is an opportunity to refine prevention strategies and develop more personalized treatment regimens.

## RESULTS

### Study population

For this retrospective study, we identified cases with large DCIS lesions (≥5 cm) and at least one of the following high-risk features: young age (<45 years), high nuclear grade, hormone receptor (HR) negative, HER2 positive, the presence of comedonecrosis, or a palpable mass. In total, 61 cases were identified for this study from the two participating sites. All participants were treated with standard of care for DCIS. **Supplementary Figure S1** provides an overview of the clinicopathologic characteristics of this cohort. The median age at diagnosis was 45 years (range: 30-88) with a median follow-up time of 8 years. The majority of patients (64%) were diagnosed with high-grade DCIS, and most cases were associated with comedonecrosis (77%). Sixty-two percent of cases were hormone receptor (HR) positive and 57% were HER2+. Lesions were subtyped as HR-positive/HER2-positive (n=21; 34%), HR-positive/HER2-negative (n=17; 28%), HR-negative/HER2-positive (n=14; 23%) or triple negative (n=9; 15%). There were 14 recurrence events in this cohort consisting of 6 DCIS and 8 invasive breast cancers.

### Biomarker assay platforms

We utilized three assay platforms to characterize the DCIS lesions and their immune microenvironments (**Figure 1**). First, Agilent 44K expression array data were generated from 52 of the 61 cases (85%). Twenty-eight gene expression signatures identifying immune cell populations, immune checkpoint genes, or immune signaling pathways were evaluated (**Supplementary Table S2)**. Second, genomic copy number data and whole exome sequencing data were generated from 34 (56%) and 17 (29%) of cases, respectively. Finally, single-plex IHC markers and mIF panels IP1.2, IP2.2, and SP1.1 were successfully performed on 54 (89%), 53 (87%), 57 (93%), and 59 (97%) of cases, respectively (**Figure 1**). Fifty-two biomarkers (immune or tumor cell populations, immune: immune cell ratios, immune checkpoint scores) were identified and evaluated from these staining panels (**Supplementary Table S3**).

**Figure 1.**
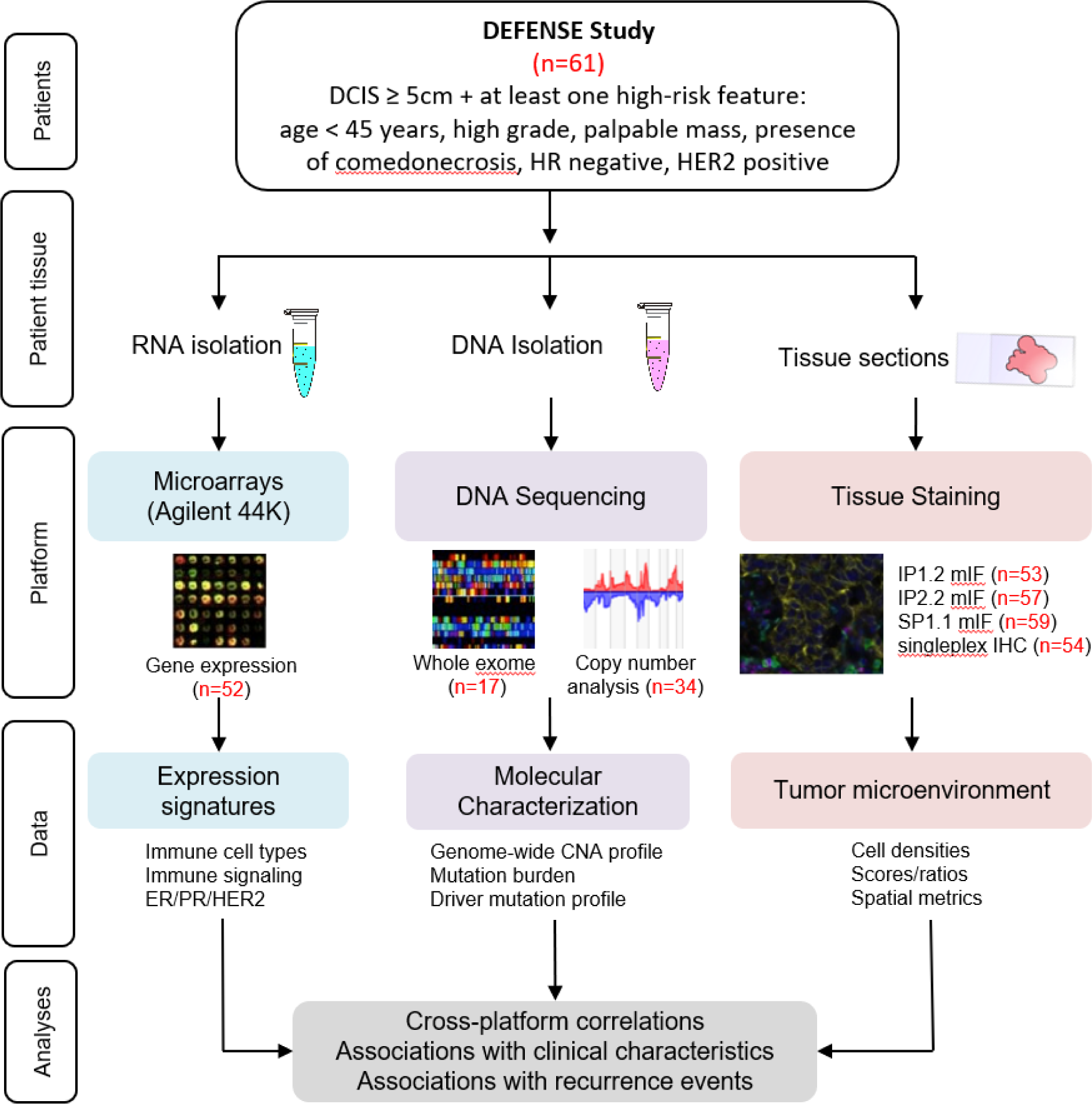
Overview of tissue collection and analyses. DCIS biopsies were processed for biomarker evaluation via gene expression microarrays, DNA sequencing, and staining tissue sections. The number of samples analyzed under each platform are shown in red.

Critical biomarkers across more than one platform, mIF and gene signatures are highly correlated eg.. mIF and gene signatures. (**Supplementary Figure S2).** Strong positive correlations between percent tumor infiltrating lymphocytes (‘TIL,’ by mIF) and the expression signatures TIL (*p* = 1.6×10^-6^) and Module4_TcellBcell_score (*p* = 1.5×10^-11^). Percentages of T cells and B cells were also strongly positively correlated with T_cell and B_cell expression signatures (*p* = 1.5×10^-4^ and *p* = 3.1×10^-11^, respectively), as were CD8^+^ T cells (CD8pT by mIF) with the CD8_T_cell and Cytotoxic_cell signatures (*p* = 1.1×10^-4^ and *p* = 4×10^-4^, respectively). Finally, a proliferation gene signature (Module11_Prolif_score) was positively correlated with Ki67^+^ tumor cells as measured by mIF (*p* = 3.1×10^-6^). In contrast, Treg cell and macrophage percentages did not correlate with their respective gene signatures.

### The genomic landscape of DCIS and immune infiltrates

We performed whole exome sequencing (WES) to identify copy number alterations (CNAs) and mutational profiles and evaluate their associations with immune infiltrates in order to identify DCIS cell-intrinsic features that predict progression. As shown in **Figure 2A**, frequently observed copy number changes included 1q, 8q, 16p, and 17q gains and 8p, 16q, and 17p losses, consistent with observations in other DCIS datasets^19–21^. The fraction of the genome altered was not significantly different between the non-recurrent and recurrent cases (DCIS or invasive recurrences; **Figure 2B**). In addition, there was no association with the fraction of the genome altered (overall CNAs, gains only, or losses only) with any immune biomarker (**Supplementary Figure S3**). However, we did observe copy number alterations at several chromosomal regions associated with various immune cell populations (**Figure 2C and Supplementary Figure S3**). In particular, gains on chromosomes 6p, 6q, and 21q, and losses on chromosomes 8p, 9p, and 17p were associated with higher TIL, T cell, CD8^+^ T cell, CD8^-^ T cell, Treg, and/or B cell infiltrates. A cytotoxic cell gene signature and two natural killer (NK) cell gene signatures were also associated with 8p chromosomal losses.

**Figure 2.**
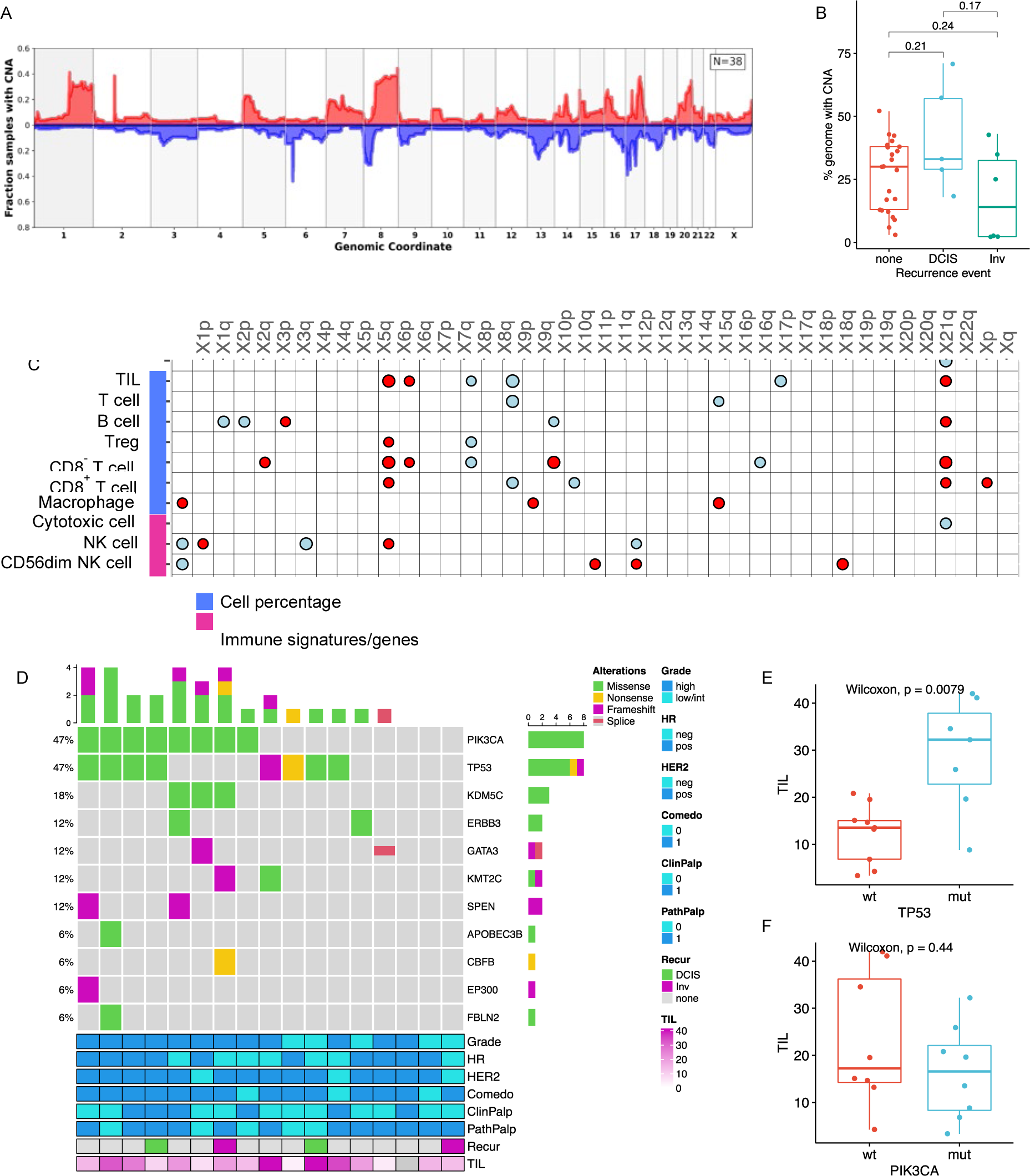
DCIS genomic landscape. (**A**). Smoothed frequency (y-axis) of CNA gains (top) and losses (bottom) smoothed along the genome (x-axis) for DCIS. (**B**). Boxplot comparing percent of genome altered by copy number in DCIS cases with none, DCIS, or invasive recurrences (Wilcoxon test p values shown). (**C**) Dot matrix plot showing associations of select immune biomarkers with chromosomal gains (red) or losses (blue). Size of dot is proportional to significance (larger dot → smaller p-value). Dots are only shown for significant associations (p<0.05; Kruskal-Wallis test). (**D**) Oncoprint diagram displaying the mutational status of driver genes commonly altered in breast cancer. Clinical characteristics and percent TILs are indicated in the bottom annotations. (**E, F**) Boxplots showing associations of TILs with TP53 and PIK3CA mutations.

Focusing specifically on breast cancer-related driver genes, the most commonly altered gene with respect to copy number status was the *ERBB2* gene (53% of cases had *ERBB2* gains) (**Supplementary Figure S4**). Other common gains (in over 40% of cases) included the *MYC*, *PPM1D*, and *NBN* loci. Common losses (in over 40% of cases) were observed for *MAP2K4*, *NCOR1*, *TP53*, *GPS2*, and *FAM86B1*. Losses were also noted in genes associated with DNA repair/genome instability (*BRCA1*, *BRCA2*, *TP53, ATRX, ZMYM3*) and cell-cycle regulation (*RB1*). Interestingly, copy number loss of *ATRX* and *ZMYM3*, genes that protect against chromosomal instability, and copy number loss of *CDKN2A*, a tumor suppressor/negative regulator in the CDK4/Rb pathway, were associated with higher densities of TILs, T cells, CD8^+^ T cells, CD8^-^ T cells, Treg cells, and/or B cells (**Supplementary Figure S5**).

The mutational status of driver genes commonly altered in breast cancer is illustrated in **Figure 2D** for 17 of the DCIS cases. *PIK3CA* and *TP53* were the most frequently mutated genes. Mutations in the *TP53* gene were significantly associated with high TIL infiltrates (**Figure 2E**) and other immune cell populations (**Supplementary Figure S6**). In contrast, mutations in *PIK3CA* were not associated with any immune cell populations (**Figure 2F** and **Supplementary Figure S6**).

### Immune cell populations are associated with DCIS clinical characteristics and outcomes

Single-plex IHC and mIF panels measured 46 immune or tumor cell populations, 4 immune:immune cell ratios, and 2 immune checkpoint scores (**Supplementary Table 2**). Immune cell densities varied across tumors. The fraction of TILs (CD3^+^ T cells plus CD20^+^ B cells) ranged from 3-59% (median 18%). As shown in **Figure 3A**, several immune cell populations identified by mIF were significantly associated with HR-negative and/or HER2-positive lesions. These included TILs, T cells, CD8^+^ T cells, Treg cells, and B cells. Proliferating cells (Ki67^+^ T cells, B cells, Tregs) and HLA-DR-positive T and B cells were negatively associated with HR status and positively associated with HER2 status. Several PD-1/PD-L1 populations were also associated with HR negativity (PD-L1^+^ immune cells, CPS score) and/or HER2 positivity (PD-1^+^ T cells, PD1^+^CD8^+^ T cells, PD-L1^+^ immune cells, CPS score). The presence of comedonecrosis was positively associated with PD1^+^CD8^+^ T cells, any PD-L1^+^ cell (tumor or immune), PD-L1^+^ immune cells, and the higher Treg:T cell ratio, and negatively associated with the overall density (cells/mm^2^) of CD68^+^ cells and the density of CD68^+^ cells within the DCIS containing ducts. Palpable lesions were positively associated with the overall density (cells/mm^2^) of CD68^+^ cells, p63^+^ cells, and CD34^+^CD68^+^ cells as well as with the density of CD34^+^ cells and CD68^+^ cells in the stroma. Finally, from the single-plex IHC markers analyzed, BAG1^+^ tumor cells were positively associated with HR status and the IHC4 score (ER/PR/HER2/Ki67) was positively associated with grade and HER2 status, and negatively associated with HR status (**Figure 3A**).

**Figure 3.**
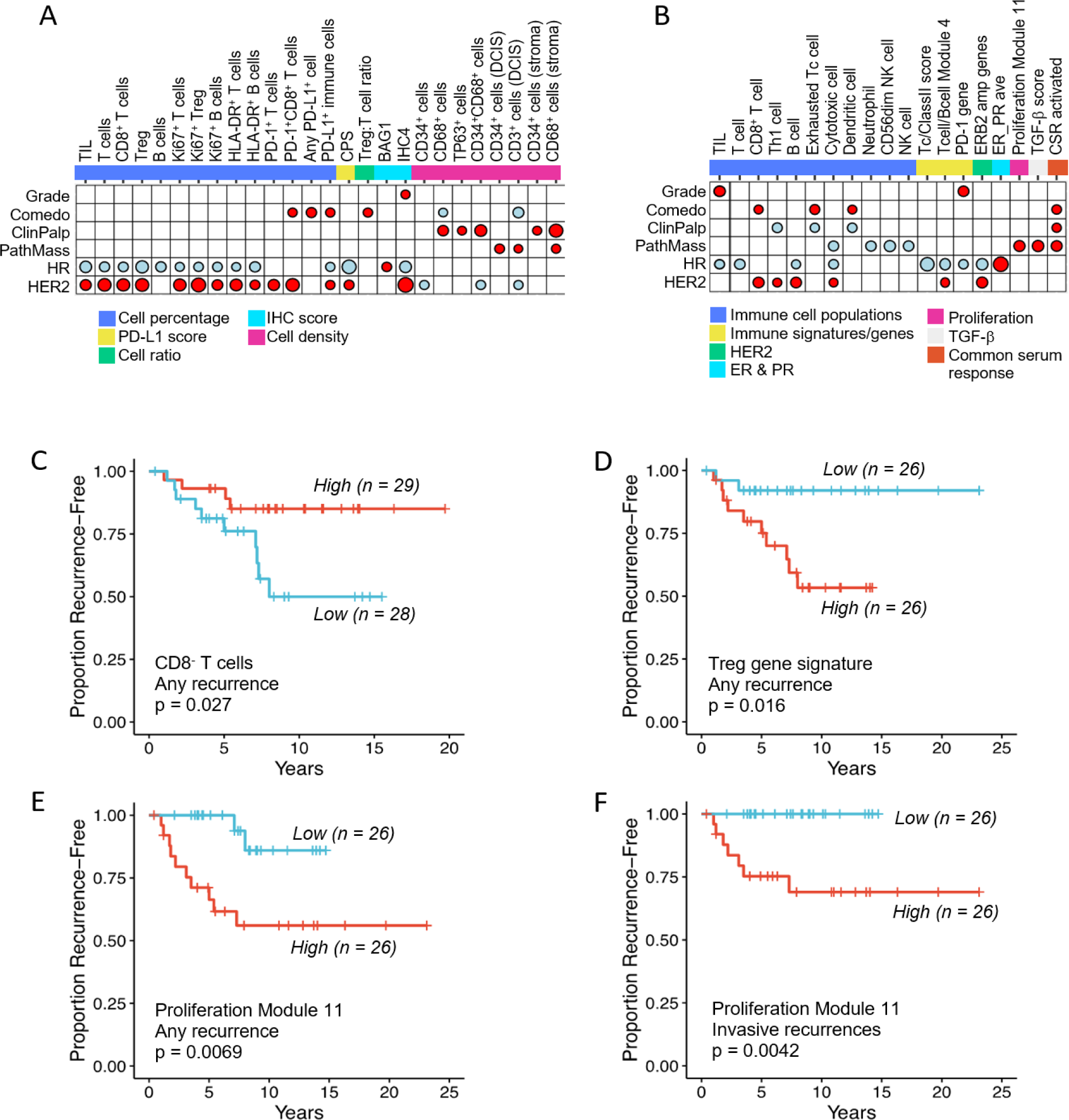
Associations of biomarkers with clinical characteristics and outcomes of high-risk DCIS. Association dot-matrix showing the level and direction of association between clinical parameters and (**A**) immune or stromal biomarkers identified by multiplex IF or single-plex IHC and (**B**) gene expression signatures. Only those biomarkers that were significant (p<0.05) with at least one clinical variable are shown. Color of dot indicates direction of association (red – positive association; blue – negative association). Size of dot is proportional to significance (larger dot → smaller p-value); Kruskal-Wallis tests used for Grade comparisons; Wilcoxon tests used for Comedo, ClinPalp, PathMass, HR, and HER2 comparisons. **C-F**. Kaplan-Meier curves showing CD8^-^ T cells were associated with good outcomes for all recurrences (**C**), a Treg gene signature was associated with poor outcomes for all recurrences (**D**), and a proliferation gene signature was associated with shorter recurrence free interval for all recurrences (**E**) and for invasive recurrences (**F**). For the dichotomization of the biomarkers, median values were used as cut points (red=high; blue=low). LogRank p values are shown.

We also evaluated 28 gene expression signatures identifying immune cell populations, immune checkpoint genes, or immune signaling pathways (**Supplemental Table S3**). Many of these expression signatures were also associated with HR negativity and HER2 positivity (**Figure 3B**). Unlike TIL and PD-1^+^ cell percentages based on mIF staining, the TIL gene signature and PD-1 gene expression were positively associated with grade.

Of the 80 mIF, single-plex IHC, and gene expression signatures examined, only 3 were associated with any recurrence event (invasive or in situ). High numbers of CD3^+^CD8^-^ T cells were associated with good outcomes (p = 0.027; **Figure 3C**) while high expression of a Treg gene signature and high expression of a proliferation gene signature were associated with poor outcomes (p = 0.016; **Figure 3D** and p = 0.0069; **Figure 3E**, respectively). When analysis was restricted to invasive recurrences **(n=8),** only the proliferation gene signature was associated with poor outcomes (p = 0.0042; **Figure 3F**).

### Spatial relationships of cells in the microenvironment of DCIS are associated with clinical characteristics and outcomes

We investigated associations between clinical characteristics/outcomes and the proximity of various immune cell populations to tumor or other immune cells. We utilized the Morisita-Horn index (MH) as a measure of the degree of colocalization of 23 different pairs of cell types, where 1 is highly colocalized and 0 means highly segregated (**Supplemental Table S3**). As shown in **Figure 4A**, the MH of 13 of these pairs were significantly associated with at least one clinical characteristic. Most of these were associated with HR negativity. The colocalizations of TIL with Treg cells (TIL_Treg.MHI), Treg cells with other T cells (Treg_Foxp3nT.MHI), and macrophages with CD3^+^CD8^-^ T cells (Mac_CD8nT.MHI) were associated with HER2-positive cases. Colocalization of T cells with B cells (T_B.MHI), CD3^+^CD8^+^ T cells with CD3^+^CD8^-^ T cells (CD8pT_CD8nT.MHI), and Treg cells with other T cells (Treg_Foxp3nT.MHI) were all positively associated with high nuclear grade DCIS.

**Figure 4.**
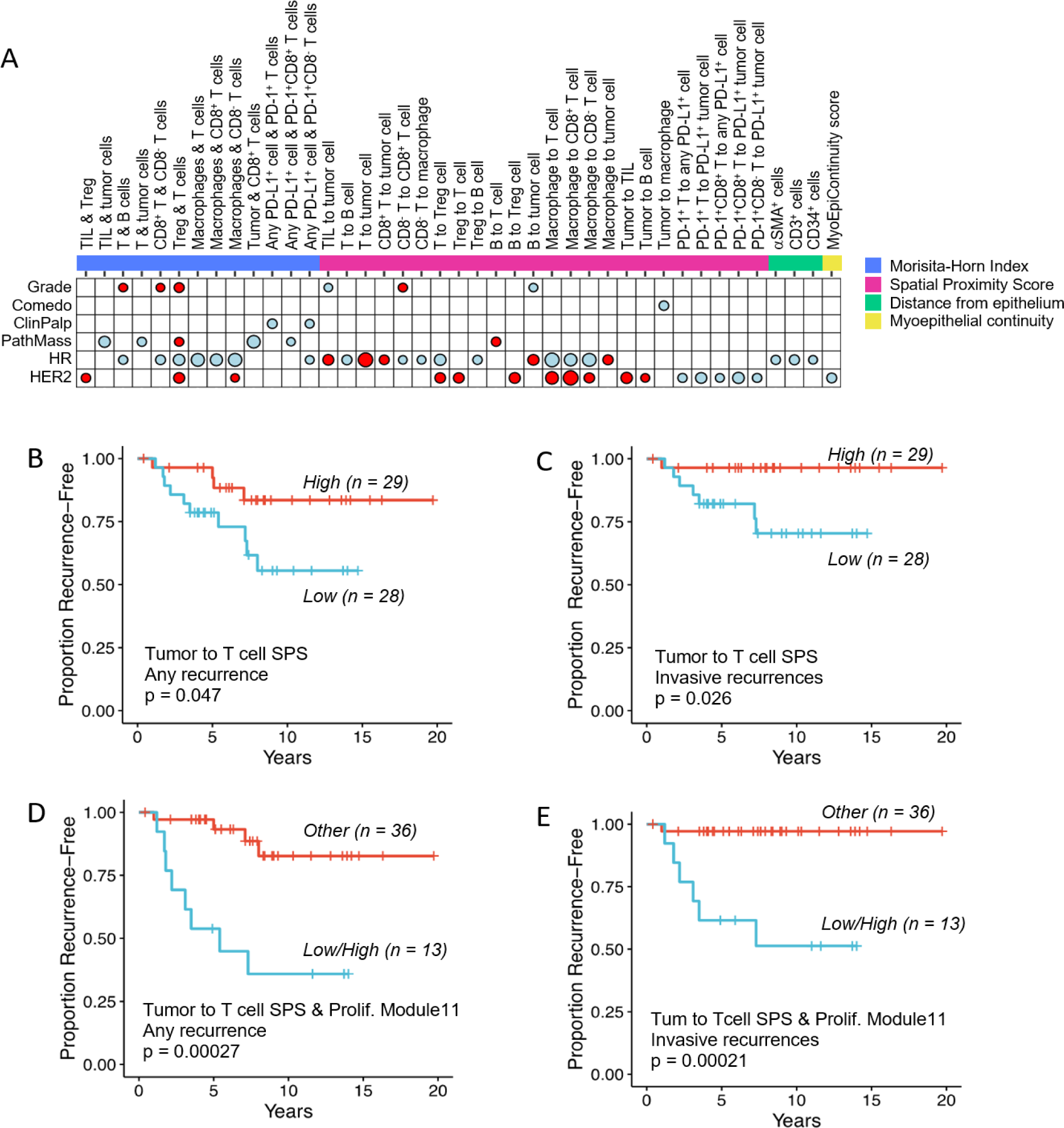
Associations of spatial biomarkers with clinical characteristics and outcomes of high-risk DCIS. **(A)** Association dot-matrix showing the level and direction of association between clinical parameters and spatial biomarkers identified by multiplex IF. Only those biomarkers that were significant (p<0.05) with at least one clinical variable are shown. Color of dot indicates direction of association (red – positive association; blue – negative association). Size of dot is proportional to significance (larger dot → smaller p-value); Kruskal-Wallis tests used for Grade comparisons; Wilcoxon tests used for Comedo, ClinPalp, PathMass, HR, and HER2 comparisons. (**B,C)** Kaplan-Meier curves showing close spatial proximity of tumor cells and T cells (high Tum to T cell SPS; red line) was associated with good outcomes for all recurrences (**B**) as well as invasive recurrences (**C**). (**D,E)** Kaplan-Meier curves showing low spatial proximity of tumor cells and T cells combined with a high proliferation score (proliferation module 11 gene signature; blue line) was significantly associated with poor outcomes for all recurrences **(D)** as well as invasive recurrences **(E)**. For the dichotomization of the biomarkers, median values were used as cut points. LogRank p values are shown.

To further characterize the spatial relationships between cells in the tumor microenvironment, we utilized a nearest neighbor distribution function, *G(r),* to evaluate the probability of a cell of type “*a*” having at least one cell of type “*b*” within a distance *r*. We defined a Spatial Proximity Score (*SPS*) as the area under the *G(r)* function curve, from 0-20μm. *SPS*s were calculated for 44 cell pairs (**Supplemental Table S3**). Twenty-four of these scores were significantly associated with at least one clinical characteristic of high-risk DCIS (**Figure 4A**). Interestingly, various *SPS*s involving PD-1^+^ cells in proximity to PD-L1^+^ cells were negatively associated with HER2 expression whereas various other cell type interactions were positively associated with HER2 expression.

We also calculated the average distances of αSMA^+^p63 ^-^, CD3^+^, CD34^+^, or CD68^+^ cells to the nearest αSMA^+^p63^+^ myoepithelial cell. Three of these markers (αSMA_DistFromEpithelium, CD3_DistFromEpithelium, CD34_DistFromEpithelium) were associated with HR negativity (**Figure 4A**).

Of the 71 spatial metrics evaluated, only the tumor to T-cell (Tum_T).SPS was associated with outcomes. A high Tum_T.SPS, indicating close proximity of tumor cells and T cells, was significantly associated with fewer invasive recurrences (p = 0.026; **Figure 4C**) or any recurrence (p = 0.047; **Figure 4B**). We next combined the Tum_T.SPS with the proliferation gene signature (Module11_Prolif_score; **Figure 2E & F)** and compared outcomes of patients with a low Tum_T.SPS and high proliferation score (n=13) to all other cases (n=36). Low spatial proximity of tumor cells and T cells combined with a high proliferation score was significantly associated with poor outcomes for all recurrences (p = 0.00027; **Figure 4D**) as well as invasive recurrences (p = 0.00021; **Figure 4E**).

### Unsupervised clustering identifies DCIS subtypes with distinct immune microenvironment profiles

Sample-wise clustering based on 94 immune cell population and cell-cell spatial measures generated on IP.1 and IP.2 revealed 6 DCIS subtypes with distinct microenvironment profiles (**Supplemental Figure S7**). One subtype (C1) only contained 1 sample; and was excluded from further consideration. Interestingly, however, this case in C1 is characterized by HR-HER2-with tumor cell PDL1 expression, low lymphocyte and macrophage measures, and microinvasion. To better characterize each DCIS subtype, we compared biomarker levels between samples within each subtype to those in other subtypes (C2-6); results were summarized in **Supplemental Table S4.** Subtype C2, comprised of HER2+ (9) and HR-HER2-(2) samples, is characterized by higher multiple immune cell populations including TIL, Tcell, Bcell, Treg, Mac, and significantly lower levels of spatial measures between PDL1p Tumor and PD1p T cell (e.g. PDL1pTum_PD1pT.SPS and PD1pT_PDL1pTum.SPS) as well as immune cell-to-tumor cell proximity scores. In contrast, samples in Subtype C3 are HR+ (1HR+HER2-, 3HR+HER2+) and have higher levels of tumor to immune cell spatial measures such as Tum_T.SPS and Tum_CD8pT MHI and lower immune to immune cell spatial measures (e.g. B_T SPS and TIL_Treg SPS). Subtype C4 were also primarily HR+ (5 HR+HER2-, 2 HR+HER2+, 1 HR-HER2-) and was characterized by lower levels of multiple immune cell populations but higher PDL1p Tumor to PD1p T cell spatial measures. Similar to C2, Subtype C5 were primarily HER2+ or HR-HER2-(10/11). However, in contrast to C2, C5 samples were characterized only by higher levels of Ki67pT, Ki69pTreg, PD1pT, PD1pCD8nT immune populations. Lastly, the largest of the subtypes (C6), which were 85% HR+ (9 HR+HER2-, 8 HR+HER2-), had lower levels Ki67p T and B cells and spatial measures between tumor and TILs (e.g. Tum_TIL.SPS and T_Tum.MHI). No significant differences in recurrence outcomes were observed between clusters.

### Myoepithelial continuity is associated with subsequent recurrence events

We derived a myoepithelial continuity score using the nearest neighbor distances between αSMA^+^p63^+^ myoepithelial cells. Myoepithelial continuity was significantly lower in DCIS compared to adjacent benign (normal or ADH) lesions (**Figure 5A-G**). Interestingly, low myoepithelial continuity was significantly associated with better outcomes for any recurrence (p = 0.038; **Figure 5H**). This suggests that low myoepithelial continuity may serve a protective function by allowing greater access of immune cells to the DCIS tumor cells. We examined correlations of various immune cell populations identified by mIF staining or gene expression signatures with the MyoEpiContinuity score and did not find any significant correlations (**Supplementary Figure S8**). We next used gene set enrichment analysis to identify pathways/ontologies that were correlated with low or high myoepithelial continuity. Low MyoEpiContinuity scores were enriched for several immune related ontologies including adaptive immune response, leukocyte proliferation, TNF superfamily cytokine production, and MHC Class II antigen assembly, processing, and presentation (**Supplementary Figure S9**). In contrast, high MyoEpiContinuity score was only enriched for one ontology related to cell adhesion. Finally, given that the presence of CD3^+^CD8^-^ T cells was associated with good outcomes for any recurrence (**Figure 3C**), we compared outcomes of patients with a low MyoEpiContinuity score plus high CD3^+^CD8^-^ T cell infiltrates (n=12) to all other cases (n=28). As shown in **Figure 5I**, a Low/High MyoEpiContinuity/CD8nT score was significantly associated with good outcomes for all recurrences (p = 0.0039).

**Figure 5.**
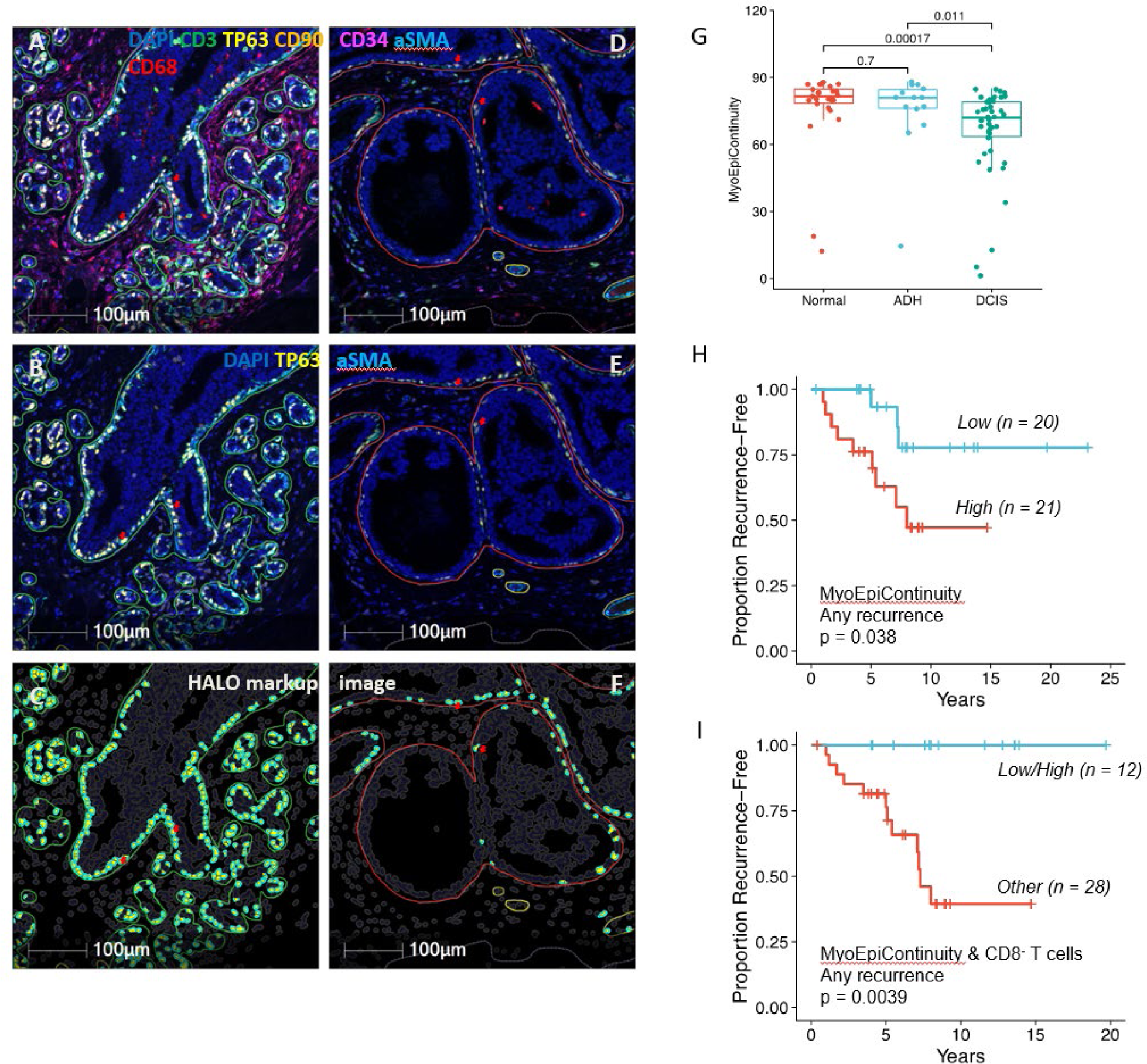
Distribution and spatial proximity of aSMA/TP63 co-expressing cells within the myoepithelium. (A-F) Representative micrographs showing distribution of aSMA/TP63 co-expressing cells within the myoepithelium of different types of lesions: (A-C) benign; (D-F) DCIS; (A,D) all markers (DAPI, CD3, TP63, CD90, CD34, aSMA, CD68); (B, E) TP63/aSMA co-expressing cells within the myepithelium; (C,F) Markup image after analysis using HALO (Indica Labs) showing only TP63/aSMA co-expressing cells. Red arrowheads showing distribution of TP63/aSMA co-positive cells between benign and DCIS lesions. (G) Boxplot comparing myoepithelial continuity in DCIS regions compared to benign (ADH) and normal regions (Wilcoxon test p values shown). (H) Kaplan-Meier curves showing low myoepithelial continuity (blue line) was associated with better outcomes for all recurrences. (I) Kaplan-Meier curves showing low myoepithelial continuity combined with high percent of CD8-T cells (blue line) was associated with good outcomes for all recurrences. For the dichotomization of the biomarkers, median values were used as cut points. LogRank p values are shown.

## DISCUSSION

This study was designed to identify genomic alterations, expression signatures, and interactions within tumor and the residents of the microenvironment, specifically immune cells, stromal elements, and the DCIS epithelium. The goal was to learn what enables large (>5cm) clinically high-risk DCIS lesions to remain as DCIS as they grow and understand what characteristics are correlated with second events. Our cohort of 61 cases with long-term follow up (median 8 years) were all treated with standard of care (surgery +/- radiation). 14 of 61 (23%) had a subsequent disease recurrence event, 8 with invasive carcinoma (13%), and 6 with recurrent DCIS (10%). These are higher recurrence rates than those reported for DCIS, as expected given the cohort design including only high-risk DCIS^6,22,23^. We performed an array of tissue-based assays and microdissections accompanied by expert (ADB) annotation of the areas of DCIS, and also a whole tissue section (bulk) transcriptome analysis. Cross-platform correlations provide new insight into the underlying biologic interactions that appear to constrain or permit progression and may reflect patterns of immunosurveillance/recognition of these lesions. Even with this more homogeneous group of large lesions, we found surprising heterogeneity.

CNAs and mutations of breast cancer driver genes, most notably TP53, were found to be significantly associated with dense immune cell infiltrates. This is similar to previously reported observations in DCIS and invasive breast cancer^14,24^. Frequently observed CNAs in our cohort included gains on chromosomes 1q, 8q, 16p, and 17q and losses on chromosomes 8p, 16q, and 17p. Breast cancer is considered to be a copy number class tumor, and these CNAs in DCIS lesions are consistent with previously published reports^19–21,25,26^. While we did not find that the fraction of the DCIS genome altered correlated with immune infiltrates or recurrence events, CNAs at certain chromosomal regions were associated with various immune cell populations. Gains on chromosomes 6p, 6q, and 21q or losses on chromosomes 8p, 9p, and 17p were associated with higher TIL, T cell, CD3^+^CD8^+^ T cell, CD3^+^CD8^-^ T cell, Treg, or B cell infiltrates. ERBB2 was the most prevalent gene with CNAs (53% of cases), which is consistent with a recent analysis of CNAs in DCIS^26^. However, our data agree with the findings of this study in that CNA at ERBB2 was not associated with recurrence events. We also found that copy number losses were common in various known breast cancer driver genes, including TP53 (>40% of cases), BRCA1, BRCA2, ATRX, and ZMYM3. Loss of TP53 function leads to overall genomic instability and resistance to DNA damage-mediated apoptosis. ATRX functions to regulate the DNA damage response and ameliorates chromosomal instability^27^. ZMYM3 deficiency results in impaired homologous recombination repair and genome instability^28^. We observed higher immune infiltrates (TILs, T cells, CD8^+^ T cells, Treg, and B cells) in DCIS harboring TP53 mutations or losses of ATRX or ZMYM3, which is consistent with the concept that genomic instability may lead to the generation of neoantigens that can activate the immune system^29^, which could potentially prevent invasion and keep the lesion contained as DCIS.

In analyzing the presence, density, and spatial relationships of various immune cells within the tumor immune microenvironment, we identified significant associations with certain clinical characteristics and outcomes. Like their invasive counterparts, we found that HR-negative and/or HER2-positive DCIS was associated with high densities of TILs, T cells, CD3^+^CD8^+^ T cells, CD3^+^CD8^-^ T cells, Treg cells, and B cells as well as proliferating (Ki67^+^) subgroups of T cells, Treg, and B cells. Several PD-1/PD-L1 immune cell populations were also associated with HR-negative or HER2^+^ DCIS. Gene expression signatures identifying immune cell populations or signaling pathways were associated with HR-negative or HER2^+^ DCIS as well. These associations are consistent with several published studies^15,30–34^; however, a recent study by Almekinders et al. that examined the relationship of DCIS immune infiltrates with recurrence events failed to find an association^30^. It should be noted that the cohort in the Almekinders study differed meaningfully from our own, in that the size of >60% of the cases in their cohort was unknown compared to the requirement that our cases all be at least 5cm in size. Furthermore, more than half of our cases (57%) were HER2^+^ compared to <30% of their cases^30^. Our cohort represents a higher risk population with overrepresentation of intrinsic subtypes known to possess a more immunologically active tumor microenvironment.

In examining 80 possible biomarkers derived from mIF, single-plex IHC, and gene expression assays, we were able to identify three that were associated with recurrence events. We found that a high density of CD3^+^CD8^-^ T cells was significantly associated with a decrease in any recurrence event. High expression of a Treg gene signature and high expression of a proliferation gene signature were each associated with greater risk of overall recurrence; the proliferation gene signature was also associated with greater risk specifically for invasive recurrence. Similar associations of a Treg gene signature and higher proportion of Treg cells in the tumor immune infiltrate with increased recurrence has been observed in the setting of early stage (Ia and Ib) non-small cell lung cancer and non-metastatic renal cell carcinoma^35,36^. Furthermore, we found that close proximity of T cells to tumor cells was significantly associated with better outcomes, with respect to any recurrence or invasive recurrences. These findings suggest that it is not solely the presence of T cells in the tumor immune microenvironment but their spatial relationship to tumor cells that predicts clinical outcomes. This is similar to findings in invasive tumors where the same subtypes i.e., HR-negative and/or HER2-positive are the most likely to have immune infiltrates and are likely to respond to immunotherapy^37^. It may also provide us with clues to what can keep tumors contained, where despite their size they remain DCIS, at least for a time. The lesions with strong immune infiltrates may be more amenable to treatment with immune cell stimulation, as we have shown^38^.

The stromal compartment appears to plays a critical immunoregulatory role and is a critical part of the story. We used spatial proximity of αSMA^+^p63^+^ myoepithelial cells to calculate a myoepithelial continuity score. Consistent with recently published work^17^, we also found that myoepithelial continuity is significantly lower in DCIS compared to benign (normal or ADH) lesions and that higher myoepithelial continuity among DCIS lesions is significantly associated with a higher recurrence risk. In addition, low myoepithelial continuity was associated with a variety of immune related pathways. These findings raise an intriguing hypothesis that myoepithelial cells may limit host immune recognition of the DCIS cells harboring neoantigens. Indeed, HER2 amplification/overexpression itself may be one of these neoantigens—with high prevalence in our cohort, but also high prevalence in all DCIS compared to lower prevalence in invasive carcinomas^39^.

Aggregating all of this complex information, we identified patterns of immune microenvironment markers expression among DCIS that help to explain the associations, shown in the **Supplementary Figure 7**. Of note, although we observed that HR-negative as well as HER2^+^ DCIS were associated with high densities of multiple immune cell populations (e.g. TILs, Tcells, Bcells), our unsupervised clustering analysis identified subsets of primarily HER2+ and HR-HER2-samples (Subtype C5) with relatively lower levels of these immune cell type markers. Unlike their Subtype C2 counterparts, which were also HER2+ or HR-HER2- and had high levels of multiple immune cell population, C5 samples exhibited higher levels of PD1pT and spatial measures of PDL1p immune cells to PD1p T cells. This analysis illustrates the heterogeneity in tumor immune microenvironment within high-risk DCIS both across and within HR/HER2 defined subtypes; and suggests classification of high risk DCIS by their microenvironment profile may offer insights beyond evaluation of single markers alone. For example, HER2+ DCIS with a strong immune infiltrate where PD1 T cells are in close proximity to PDL-1+ cells may indicate an environment where ongoing immune activity is preventing invasive recurrence. These insights are the basis for initiating immunotherapy trials for DCIS^30,38–41^.

In summary, this unique DEFENSE study of large DCIS that Nevertheless Stay Encapsulated demonstrates that DCIS can be stratified based upon features of the tumor immune microenvironment which, in turn, correlates with intrinsic epithelial subtypes including propensity for high neo-antigenicity, along with myoepithelial continuity and finally immune activation. It is the interplay among these features that determine the fate of these lesions. The observations we describe have the potential to guide clinical decision-making including the safety of active surveillance (watchful waiting) for lower risk lesions as well as improved therapy including immunomodulation that can be targeted to higher risk lesions which are most likely to be responsive and permit trafficking of immune cells past the myoepithelial layer to the lesions within the ducts.

## METHODS

### Study population and clinical data collection

We identified cases with large DCIS lesions (≥ 5 cm on imaging or pathology) and at least one of the following high-risk features: young age (<45 years), high nuclear grade, hormone receptor negative, HER2 positive, the presence of comedonecrosis, or a palpable mass. Retrospective clinical characteristics were obtained through chart review without direct patient contact. **Supplementary Figure S1** summarizes the data for the patients in this cohort. All patients enrolled on this study provided written consent for their tissue to be used for research. The Institutional Review Board (IRB) of the University of California, San Francisco (UCSF) approved the conduct of this study with other University of California sites and the University of Vermont (UVM) listed as collaborating sites under protocol #16-20550.

### Single-plex immunohistochemistry

Immunohistochemistry (IHC) was performed as previously reported^42^. Formalin fixed paraffin embedded (FFPE) tissue sections (4 μm) were prepared on Superfrost Plus microscope slides (Thermo Fisher Scientific). Sections were deparaffinized followed by antigen retrieval in CC1 buffer (pH 9, 95°C; Roche), endogenous peroxidase blocking, and then incubation with primary antibodies. Chromogenic detection was performed with VECTASTAIN Elite ABC-HRP kit (Vector Laboratories) followed by counterstaining with hematoxylin. Images were recorded with the Aperio Leica ScanScope XT (Leica Biosystems, Richmond, IL) using x20 objective lens. Primary antibodies used for this study are listed in **Supplementary Table S1**.

### Multiplex immunofluorescence analysis with multispectral imaging

Multiplex immunofluorescence (mIF) analysis was performed with tyramide signal amplification using Opal fluorescent dyes (Akoya Biosciences). Three mIF panels were used in this study (IP1.2, IP2.2, and SP1.1). IP1.2 consisted of antibodies against CD3, CD20, Foxp3, HLA-DR, Ki67, and pancytokeratin AE1/AE3. IP2.2 consisted of antibodies against CD3, CD8, CD68, PD-1, PD-L1, and pancytokeratin AE1/AE3. SP1.1 consisted of antibodies against CD3, CD34, CD68, CD90, αSMA, and p63. Staining was performed with a BondRX autostainer (Leica Biosystems). Antibody/Opal dye pairings, concentrations, and order in the panel were optimized as previously reported^43^. Antibody clones, sources, and pairings with Opal dyes are listed in **Supplementary Table S1**.

Slides stained with mIF panels IP1.2 and IP2.2 were scanned using the Vectra Polaris multispectral imaging system (Akoya Biosciences). Regions of interest were selected in Phenochart (Akoya) and spectrally unmixed using inForm software (Akoya). Unmixed image tiles were then stitched together and cell segmentation performed in QuPath^44^. Cell phenotyping was performed using flow cytometry based R packages (flowDensity, flowCore, openCyto). Twenty-one cell populations (reported as a percentage of total cell counts), 4 cell:cell ratios, and 2 immune checkpoint scores were identified with these two panels (**Supplementary Table S3**). PD-L1 staining was assessed from IP2.2 using the tumor proportion score (TPS) and the combined positive score (CPS). The TPS was defined as the number of PD-L1^+^ tumor cells divided by the total number of tumor cells multiplied by 100. The CPS was defined as the number of PD-L1^+^ tumor cells and PD-L1^+^ immune cells, divided by the total number of tumor cells multiplied by 100.

Slides stained with mIF panel SP1.1 were scanned on a Nikon A1R HD laser scanning confocal microscope equipped with a spectral detector and 405 nm, 445 nm, 488 nm, 514 nm, 561 nm, 640 nm laser lines (Nikon Instruments Inc., Melville NY). The mIF images obtained on the confocal microscope were spectrally unmixed and analyzed using HALO and HALO AI analysis packages (Indica Labs). Using a custom trained HALO AI DenseNet classifier, ROIs were classified as either tumor or stroma. The tumor regions were further designated as normal, DCIS or ADH based on a pathologist’s annotations on the H&E images. Segmentation and phenotyping of cells within these classified regions was performed with HALO module “HighPlex FL” (version 3.2.1). Seven cell populations were phenotyped with this panel and are reported as cell densities (counts/mm^2^) in the overall tissue or separately within DCIS or stromal areas (**Supplementary Table S3**).

### Analysis of spatial relationships of cells in the microenvironment of DCIS

For slides stained with IP1.2 and IP2.2 we employed spatial point pattern analyses using the R package “spatstat”^43^. To study the spatial colocalization of cancer cells and immune cells, each mIF image was virtually divided into non-overlapping hexagons (100 μm sides). The number of cancer cells and immune cells within each hexagon was counted. To calculate colocalization of two cell types, A & B, we applied the Morisita-Horn (MH) similarity index^44,45^ to the data using the R package ‘divo’. The MH index ranges from 0 (no cells of type A share the same hex with a cell of type B), indicating highly segregated cell populations, to 1 (an equal number of type A and type B cells in each hex), indicating the two cell types are highly colocalized. MH indices were calculated for 23 pairs of cell types (**Supplemental Table S3**).

To further examine spatial relationships, we utilized the nearest neighbor distribution function, *G(r),* to quantify the relative proximity of any two cell types. This function is a spatial distance distribution metric that represents the probability of finding at least one cell of a given type within a radius (*r*) of another cell type. The *G(r)* function computations were performed using the R package “spatstat”. We defined a Spatial Proximity Score (*SPS*) as the area under the *G(r)* curve (AUC) for *r* = 0 to 20 μm. A large *SPS* indicates close proximity between the cell populations analyzed. The *SPS* is directional. For example, the score for T cells within 20 μm of tumor cells, denoted *Tum_T.SPS* will be different than the score for tumor cells within 20 μm of T cells (*T_Tum.SPS)*. *SPS* scores were determined for 44 pairs of cell types (**Supplemental Table S3**). The 20 μm interval was selected to build upon prior work in the field^46–48^. To put this into perspective, a lymphocyte has a diameter of ∼10 μm.

Spatial analyses from slides stained with SP1.1 were performed with HALO Highplex FL (version 3.2.1) using the Proximity Analysis algorithm from the Spatial Analysis module within HALO. The distance between cells positive for each marker and cells that were co-positive for p63 and αSMA was measured and is reported as “marker” DistFromEpithelium. In order to quantify myoepithelial continuity, the average nearest neighbor distances between αSMA^+^/p63^+^ myoepithelial cells within the regions of DCIS were calculated. A myoepithelial continuity score was then derived by reverse scoring these distances such that a high score (high continuity) = short distance between myoepithelial cells (close proximity) and a low score (low continuity) = greater distance between these cells. Reverse scoring was performed by taking the maximum average nearest neighbor distance between myoepithelial cells, adding 1 to this distance, and then subtracting the nearest neighbor distances from this value to get the reverse scored values.

### Characterization of samples by their immune profiles

To characterize samples by their microenvironment profiles as assessed via the cell population and spatial markers, unsupervised consensus clustering of 57 samples based on 94 markers generated from IP1.1 and IP1.2 (**Supplemental Table S3**) was performed using the ConsensusClusterPlus package in R^49^. Specially, biomarkers were median-centered and scaled to a standard deviation of 1. Euclidean distances between samples were calculated based on available (pairwise complete) data for input into the clustering algorithm. Hierarchical clustering using the Ward’s minimum variance method (i.e. ward D2 inner linkage option) with 80% subsampling was performed over 1000 iterations; and the final consensus matrix was clustered using average linkage. The number of clusters (k) was selected based on the area under the empirical cumulative distribution function (CDF) curve at a point where the area reaches an appropriate maximum and further separation provides minimal relative change. A final heatmap display of the samples’ microenvironment profiles was created using the pheatmap package [Kolde R (2019). pheatmap: Pretty Heatmaps. R package version 1.0.12, <https://CRAN.R-project.org/package=pheatmap>] in R. Biomarkers characteristics of each sample cluster (vs. all others) was identified using Wilcoxon rank sum test with Benjamini Hochberg false discovery rate correction.

### Gene expression

FFPE tissue sections were sent to Agendia, Inc, for RNA extraction and gene expression on Agilent 32K expression arrays (Agendia32627_DPv1.14_SCFGplus with annotation GPL20078). MammaPrint risk classification (High vs. Low risk) and BluePrint molecular subtype (Luminal, HER2 or Basal type) were calculated. For each array, the green channel mean signal of each probeset was log2-tranformed and centered within array to its 75^th^ quantile as per the manufacturer’s data processing recommendations. All values flagged for non-conformity are NA’d out; and a fixed value of 9.5 was added to avoid negative values. Probeset level expression data was mean-collapsed to gene level by gene symbol, yielding a normalized expression dataset comprising 54 samples from 52 patients x 22584 unique genes. Twenty-eight previously published immune-related signatures, along with a cell proliferation and ER/PR signature, were computed using the scoring methods described in Supplemental Table 3 from the gene-level normalized expression data.

Gene set enrichment analysis (GSEA) was used to determine pathways enriched in DCIS with high or low myoepithelial continuity scores. GSEA was performed using the R package fgsea (v1.24.0)^25^ and the human MSigDB collection of gene sets derived from the GO biological process ontology (c5.go.bp.v2023.1.Hs.symbols.gmt). Pathways were considered enriched if adjusted p-values < 0.05.

### Whole exome and whole genome sequencing

Whole exome sequencing including copy number alterations (CNAs) and mutations was performed on 17 cases, and low-pass whole genome sequencing including CNAs was performed on 34 cases. Tissue from DCIS involved ducts was obtained by laser capture microdissection (ArcturusXM system; ThermoFisher). DNA was extracted using the QIAmp DNA Micro Kit (Qiagen). Libraries were prepared using Ultra II FS library preparation kit (New England Biolabs) and amplified with KAPA HiFi HotStart Real-time PCR using KAPA P5 and P7 primers (KAPA Biosystems).

#### Exome sequencing

Exome sequencing was performed as previously described^14,24,50^. Sequencing libraries were deconvoluted using bcl2fastq. Sequencing data were analyzed using bcbio-nextgen (v1.1.6) as a workflow manager. Adapter sequences were trimmed using atropos (v1.1.22), the trimmed reads were subsequently aligned with bwa-mem (v0.7.17) to reference genome hg19, then PCR duplicates were removed using biobambam2 (v2.0.87). Additional BAM file manipulation and collection of QC metrics were performed with picard (v2.20.4) and samtools (v1.9). Contamination was measured using verifyBamID2 (v1.0.6). Samples with less than 90% of their bases covered at 10x or contamination estimated to be >5% were not used for mutational analysis.

#### Low-coverage whole-genome sequencing

For low-coverage whole-genome sequencing, sample libraries were sequenced to a target coverage of 0.5x using paired-end 2×100bp reads on a NovaSeq 6000 (Illumina). Sequencing libraries were deconvoluted using bcl2fastq. Adapter sequences were trimmed from the raw fastq files using atropos (v1.1.31). The trimmed reads were then aligned to GRCh38 using bwa-mem (v0.7.17). Duplicate reads were then marked using biobambam (v2.0.87). Overall genome-wide coverage was measured using mosdepth (v0.2.6), and contamination was measured using verifyBamID2 (v1.0.6). Samples with less than 0.45x coverage or were estimated to be >5% contaminated were removed from downstream analyses.

#### Copy number alterations

CNAs were called using CNVkit (v0.9.9) in “wgs” mode with average bin size set at 100,000 bp. A set of unrelated normal tissues sequenced with the same protocol were used to generate a panel of normals used during CNA calling. Any bins with a log_2_ copy ratio lower than -15 were considered artifacts and removed. Breakpoints between copy number segments were determined using the circular binary segmentation algorithm (p < 10^−4^). Copy number genomic burden was computed as the sum of sizes of segments in a gain (log_2_(ratio) > 0.3) or loss (log_2_(ratio) < − 0.3) over the sum of the sizes of all segments. Gene-level CNA and exome-sequencing-based CNA were measured as previously described^50^.

#### Pathogenic germline variants

For all cases, we identified variants that were either “Pathogenic” or “Likely-Pathogenic” in ClinVar and had a variant allele frequency (VAF) of at least 15% in the following breast cancer susceptibility genes: *ATM, BARD1, BLM, BRCA1, BRCA2, BRIP1, CDH1, CHEK2, EPCAM, FAM175A, FANC, MLH1, MRE11, MSH2, MSH6, MUTYH, NBN, NBS1, NF1, PALB2, PMS2, PTEN, RAD50, RAD51C, RAD51D, RECQL, STK11, TP53, XRCC, MEN1,* or *PPM1D*.

#### Somatic mutations

Somatic mutations and removal of putative germline variants were determined from tumor-only data as previously described^50^.

### Statistical analyses

Statistical analyses were performed using R. Associations of clinical characteristics with immune cell populations, gene expression signatures, and spatial metrics are presented as dot matrix plots. Associations of immune biomarkers with chromosomal gains/losses are also presented as dot matrix plots. Grouped data box plots are presented with individual sample points. Data were compared between two groups with Wilcoxon’s test or between multiple groups using the Kruskal-Wallis test. Associations of biomarkers with subsequent recurrence events were assessed in univariate analyses using Kaplan-Meier curves and the Log Rank test statistic. Multiple testing corrections were not performed.

## ACKNOWLEDGMENTS

The authors acknowledge support from the NIH/NCI MCL Consortium grant and the HTAN PCA Pilot grant (U01CA196383 and U01CA196406). AG was funded by T32CA251070. The authors thank Jeff Matthews (Quantum Leap Health) for editorial assistance.

## Supplementary Information

**Supplementary Figure S1.**
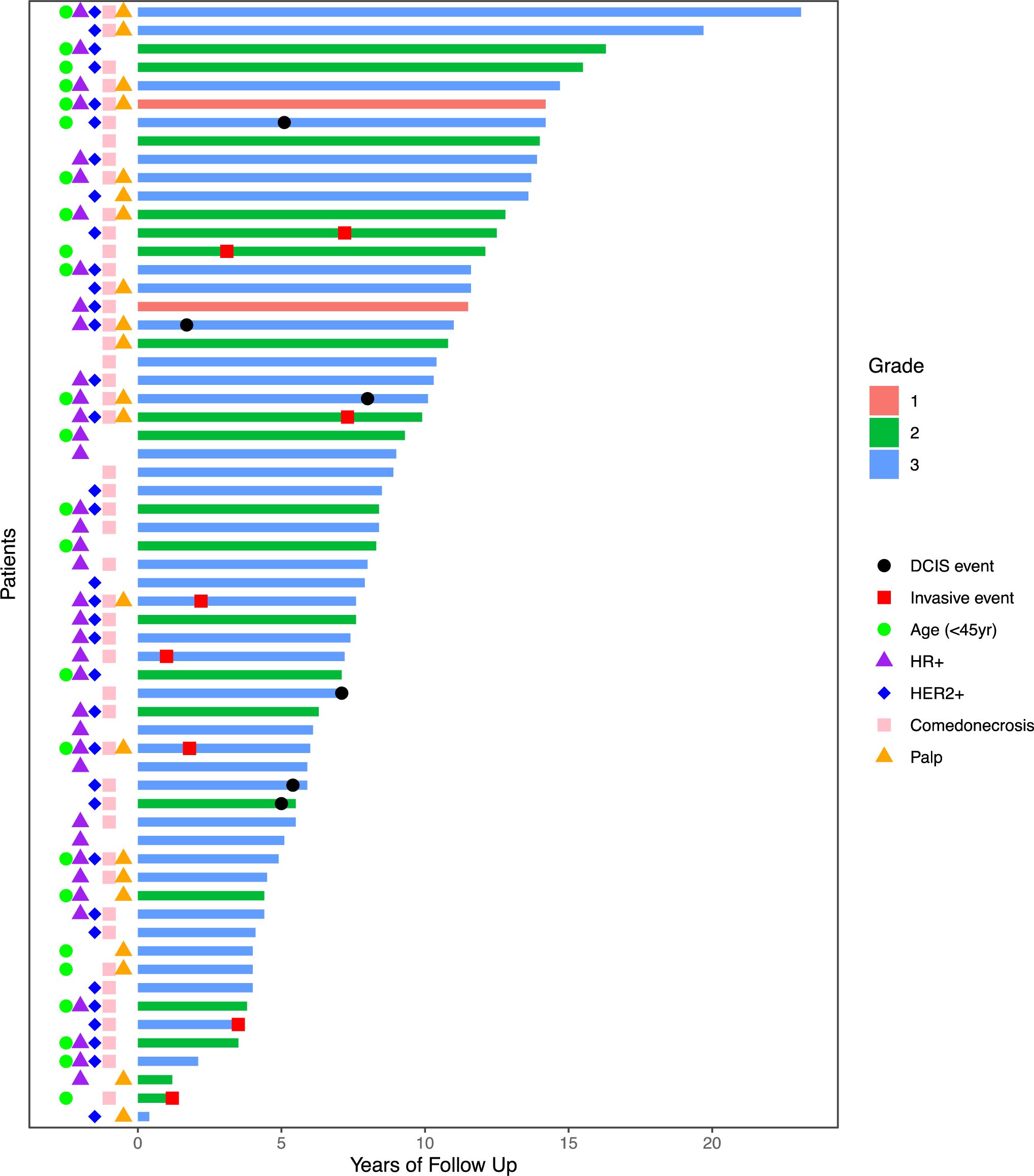
Swimmer plot showing clinical characteristics of DCIS patients in the DEFENSE cohort. Patients are ordered by years of follow-up relative to original DCIS diagnosis. Clinical characteristics are shown along the y axis. Bars are color coded according to grade. Timing of subsequent DCIS or invasive events are also denoted.

**Supplementary Table S1.** Antibodies used in mIF and IHC assays.

| **Marker** | **Clone** | **Antibody source** | **Fluorophore** |
| --- | --- | --- | --- |
| **mIF Panel IP2.2** |  |  |  |
| PD-L1; immune checkpoint | E1L3N | Cell Signaling | Opal520 |
| PD-1; immune checkpoint | EPR4877 | Abcam | Opal620 |
| CD8; cytotoxic T cell marker | 4B11 | Leica | Opal570 |
| pan-cytokeratin; epithelial cell marker | AE1/AE3 | Dako | Opal690 |
| CD68; macrophage marker | PG-M1 | Dako | AlexaFluor750 |
| CD3; T cell marker | 2GV6 | Ventana/Roche | Opal480 |
| **mIF Panel SP1.1** |  |  |  |
| CD3 | LN10 | Leica | Opal520 |
| aSMA | ASM1 | Leica | Opal650 |
| p63 | D9L7L | Cell Signaling | Opal540 |
| CD90 | EPR3133 | Abcam | Opal570 |
| CD34 | QBEND/10 | Leica | Opal620 |
| CD68 | 514H12 | Leica | Opal690 |
| **Single-plex IHC stains** |  |  |  |
| AR | SP107 | Cellmarque |  |
| BAG1 | Y166 | Abcam |  |
| COX2 | CX294 | DAKO |  |
| ER | SP1 | Abcam |  |
| PgR | pgr636 | DAKO |  |
| Ki67 | MIB-1 | DAKO |  |
| p16 | E6H4 | Ventana |  |
| p53 | DO-7 | DAKO |  |

**Supplementary Table S2.**
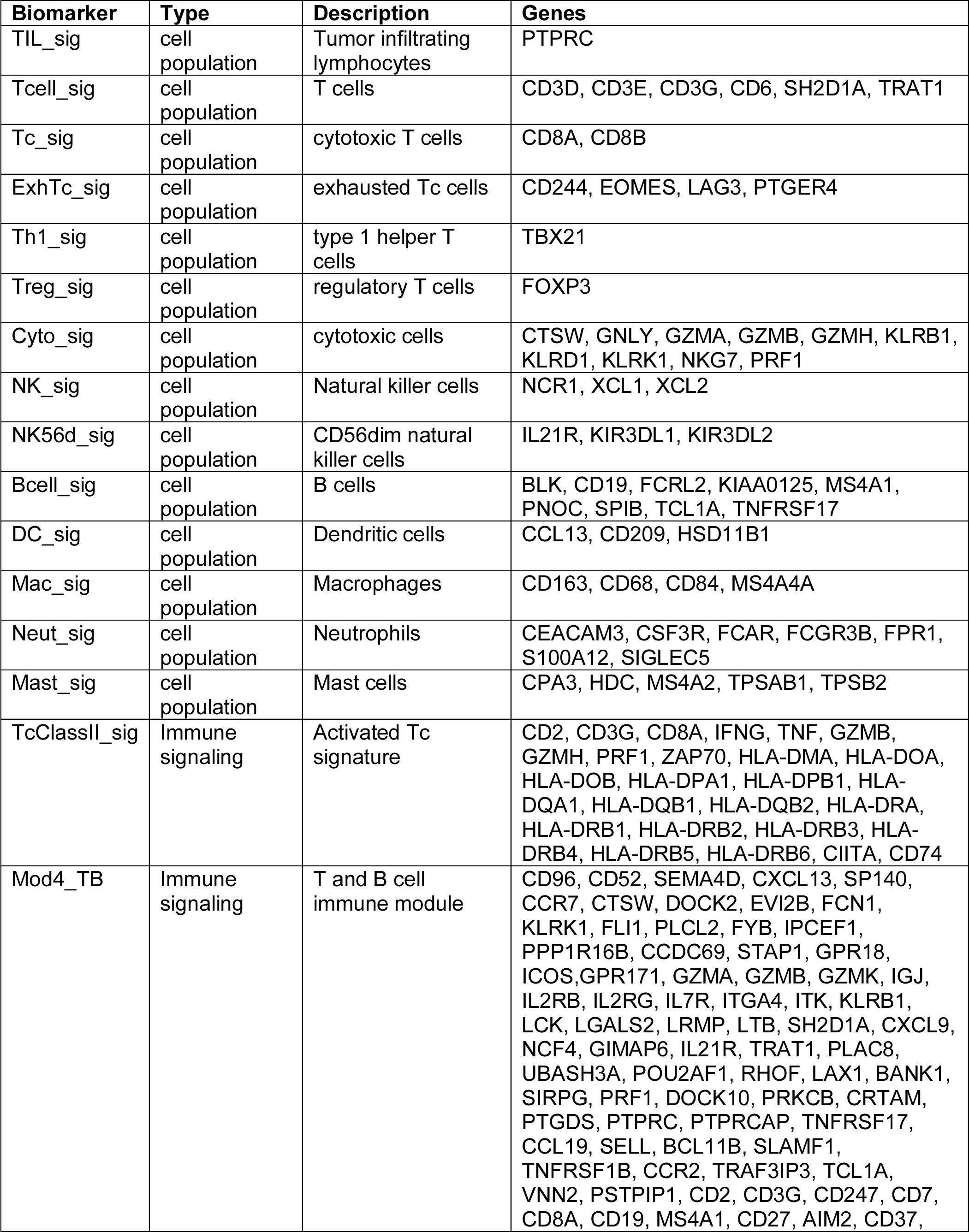
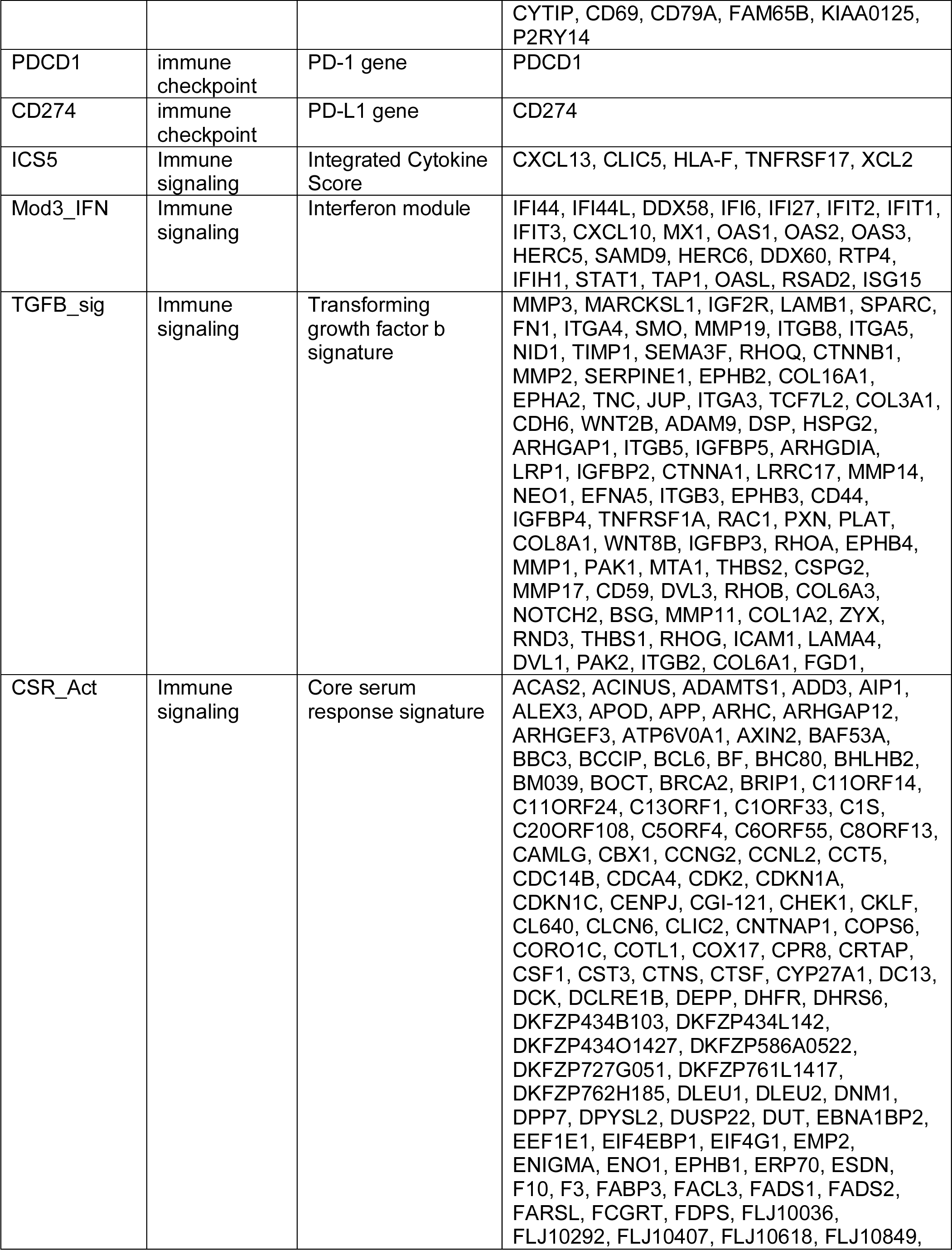
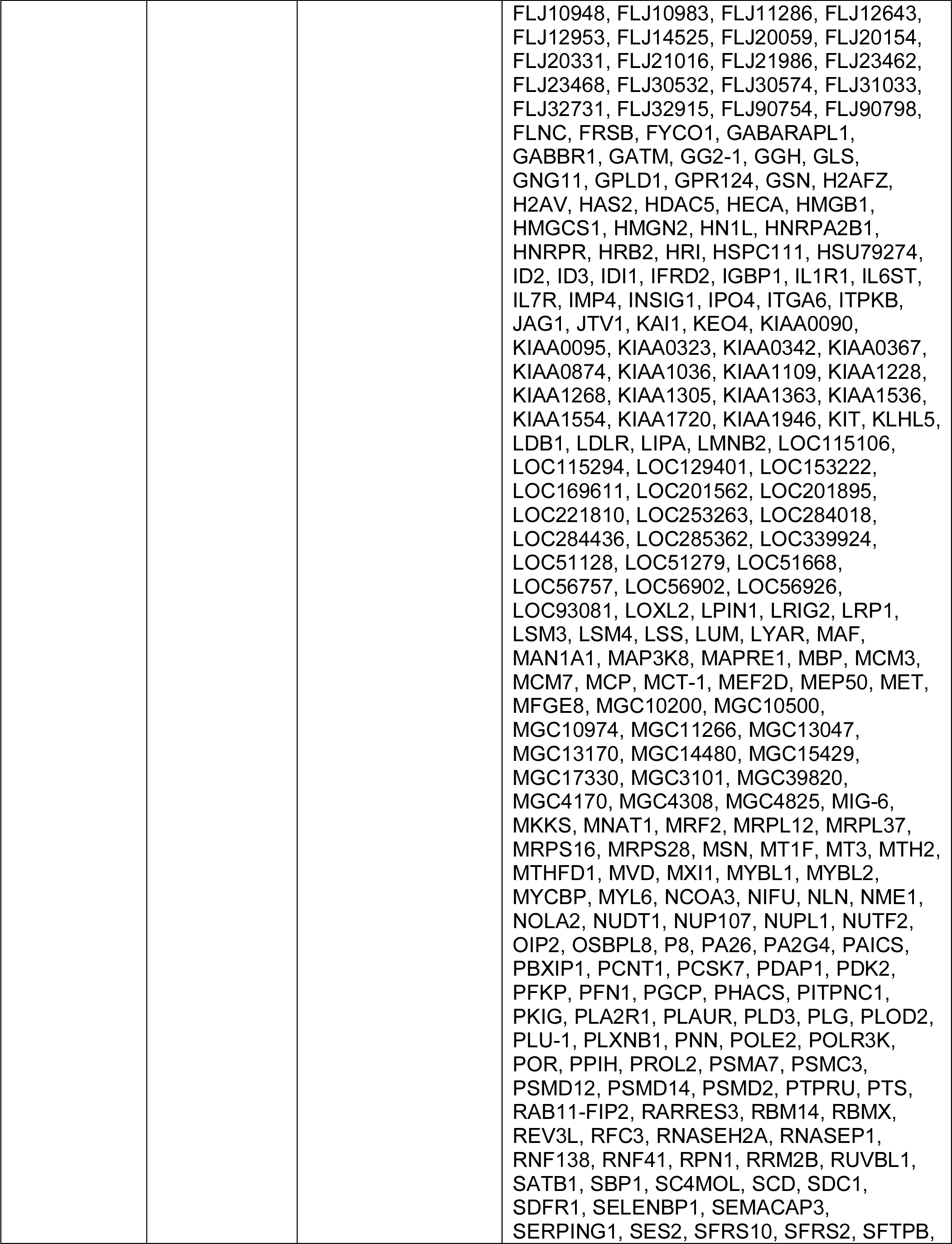
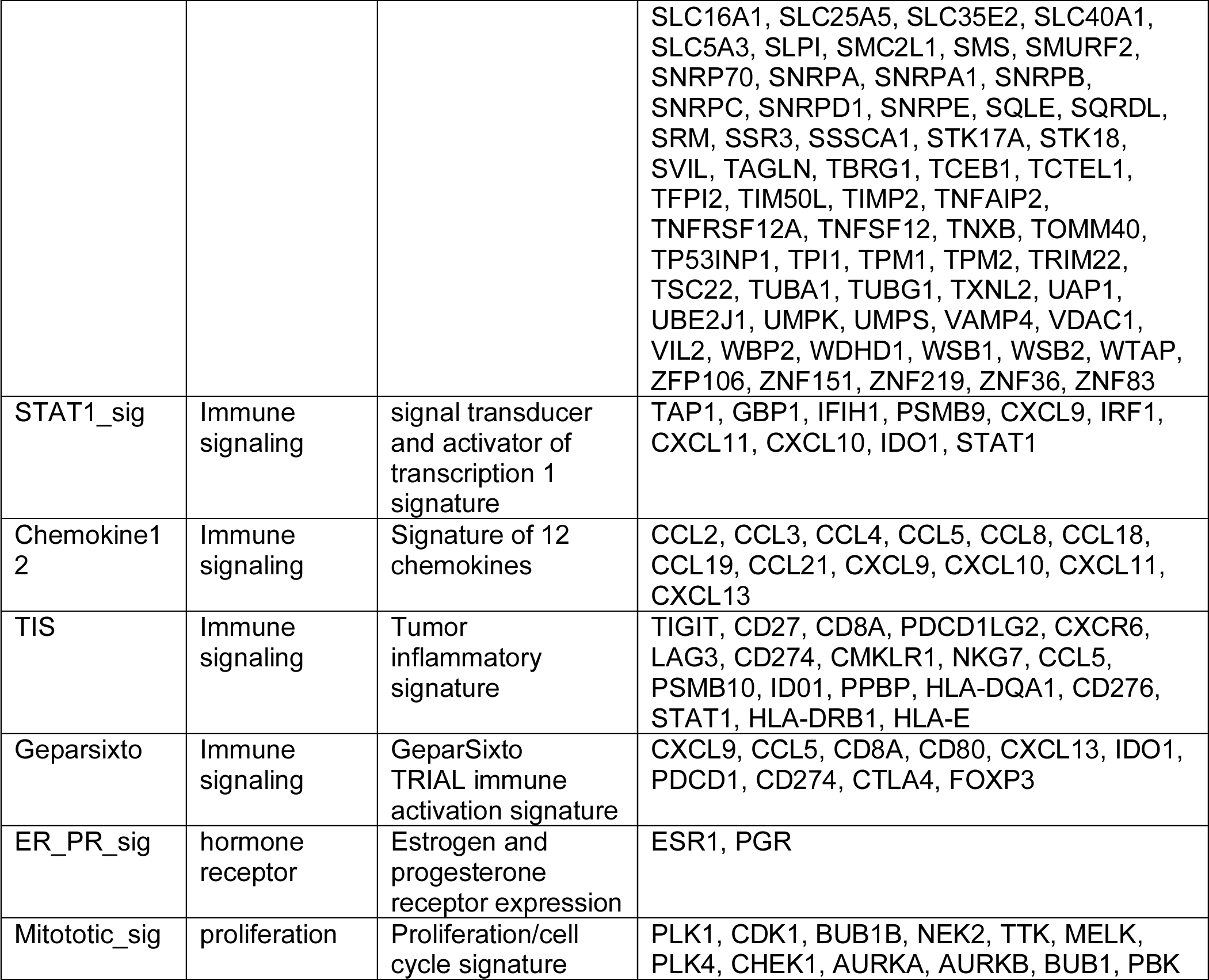
Expression array based biomarkers evaluated in this study.

**Supplementary Table S3.**
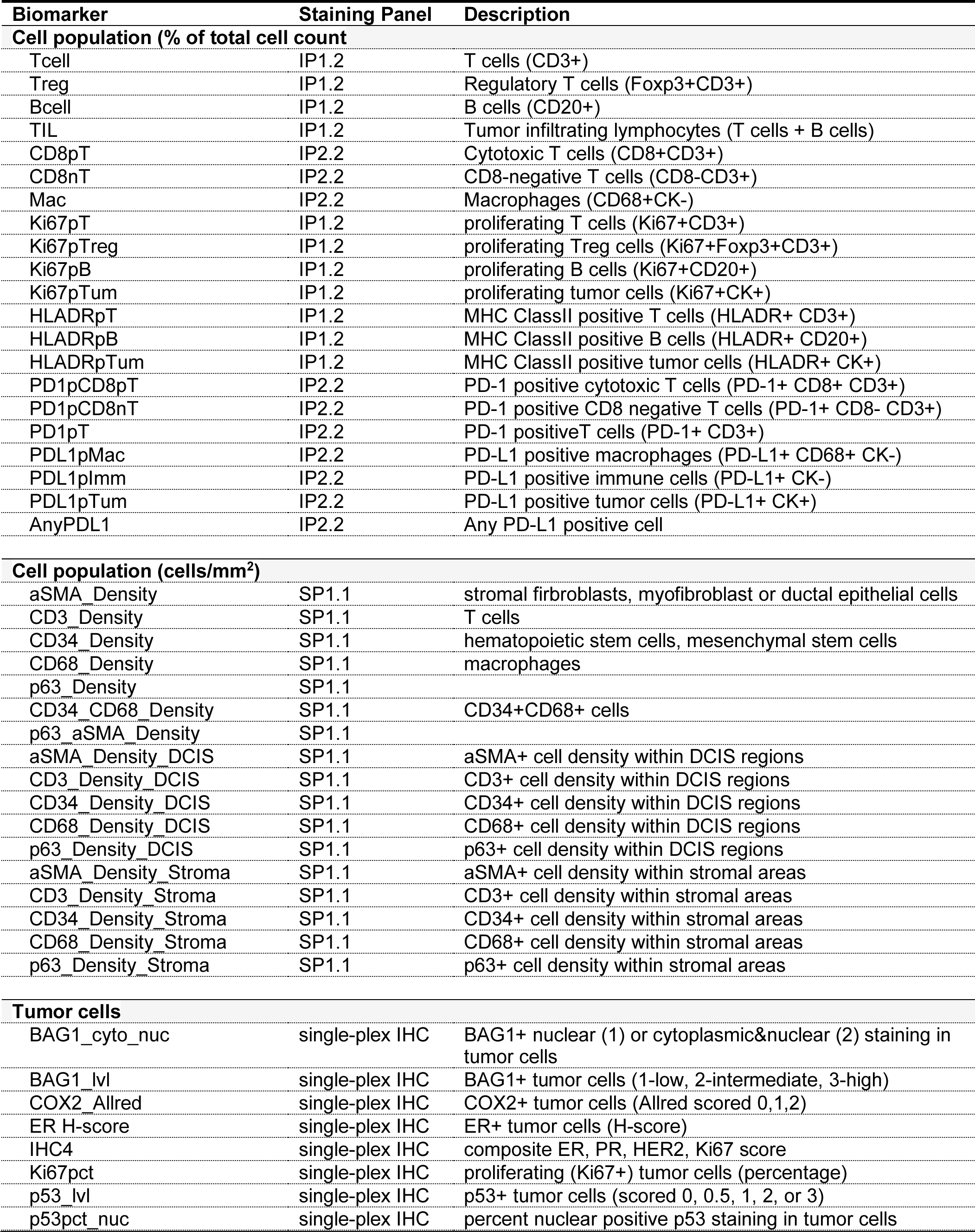
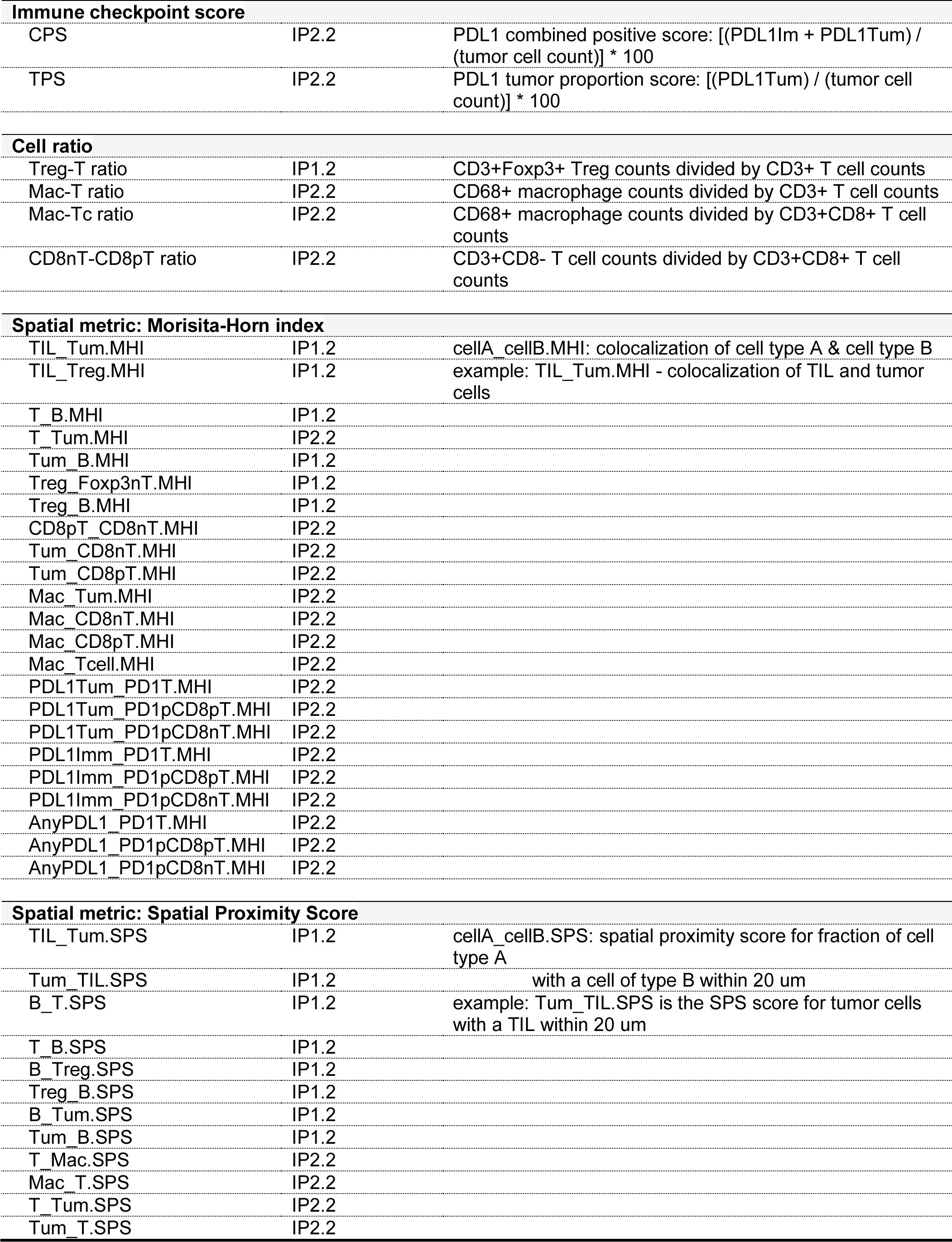
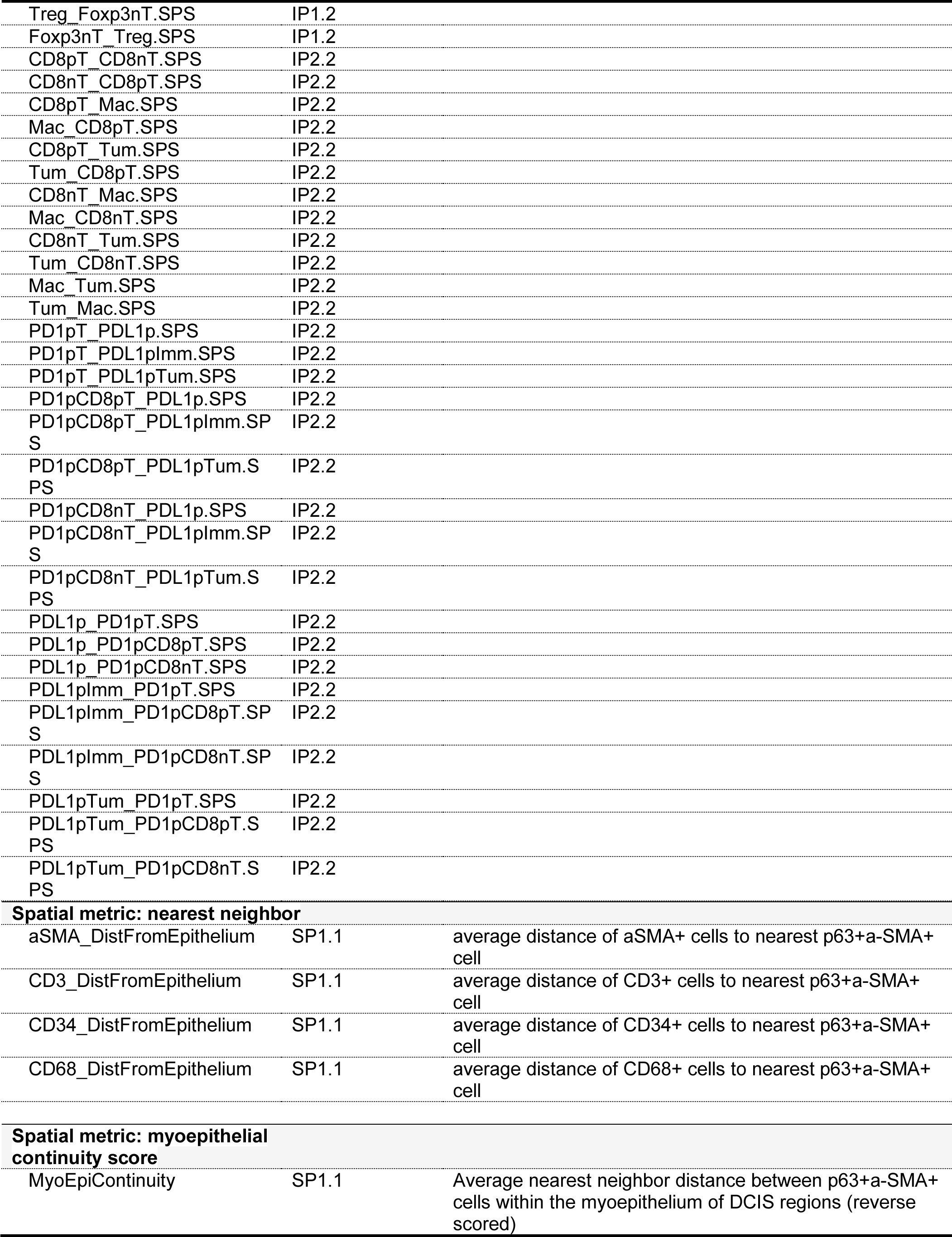
Immune biomarkers evaluated in this study via IHC and mIF.

**Supplementary Figure S2.**
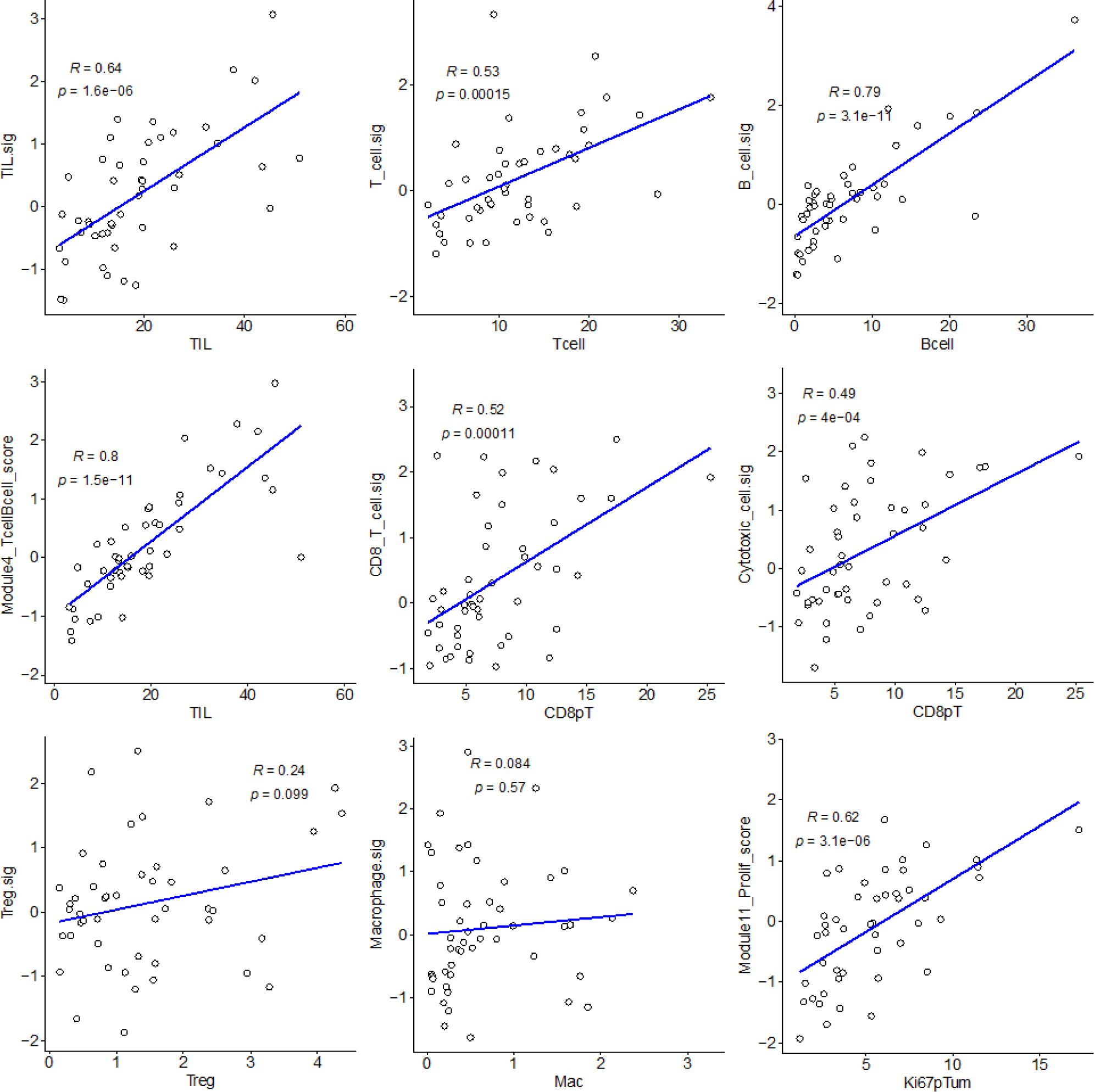
Correlations between cell populations identified by mIF staining (x-axes) and corresponding gene expression signatures (y-axes). Pearson R and p values

**Supplementary Figure S3.**
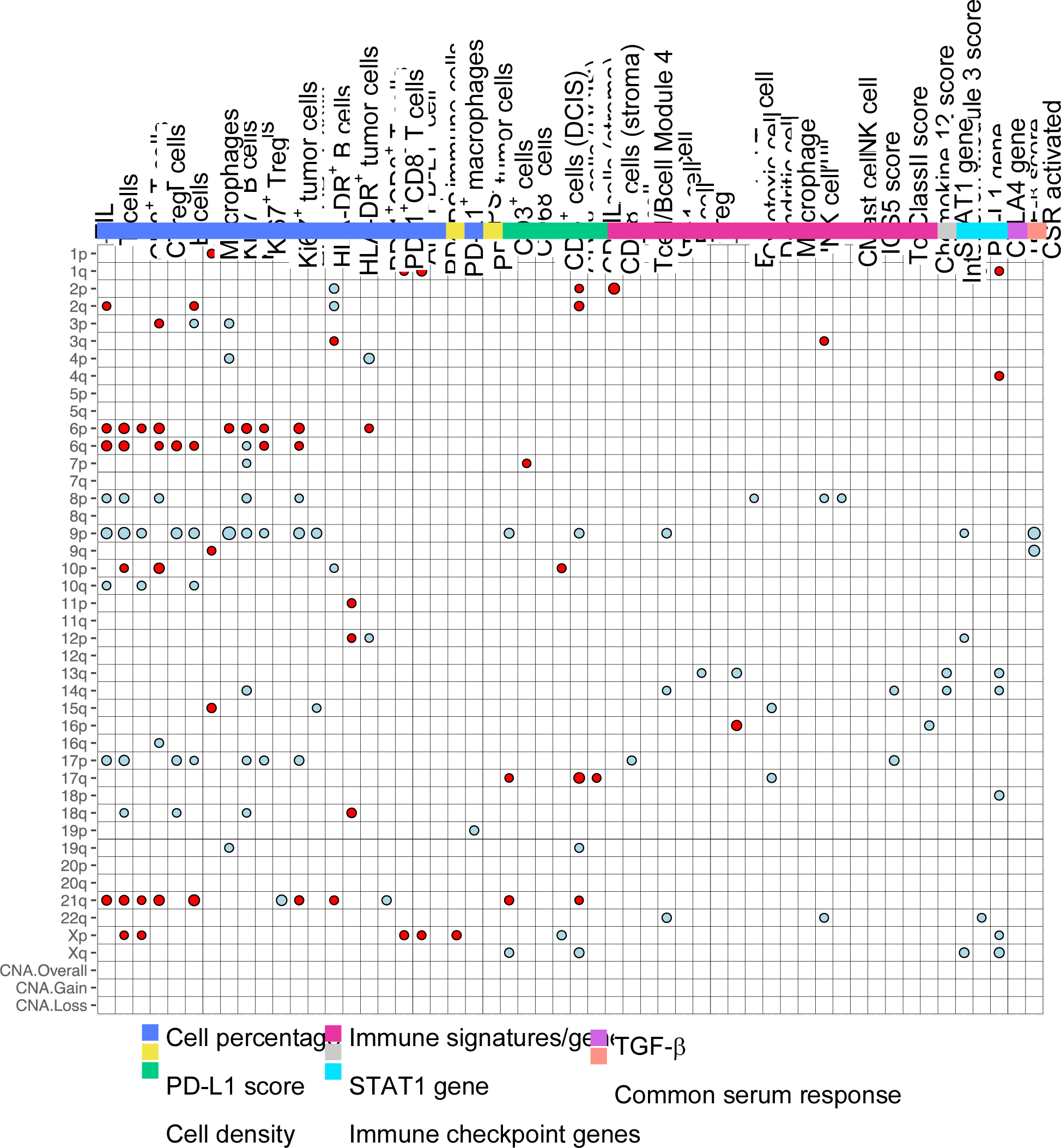
Associations of chromosomal aberrations with immune biomarkers. Color of dot indicates association with gains (red) or losses (blue). Size of dot is proportional to significance (larger dot → smaller p-value). Dots are only shown for significant associations (p<0.05; Kruskal-Wallis test).

**Supplementary Figure S4.**
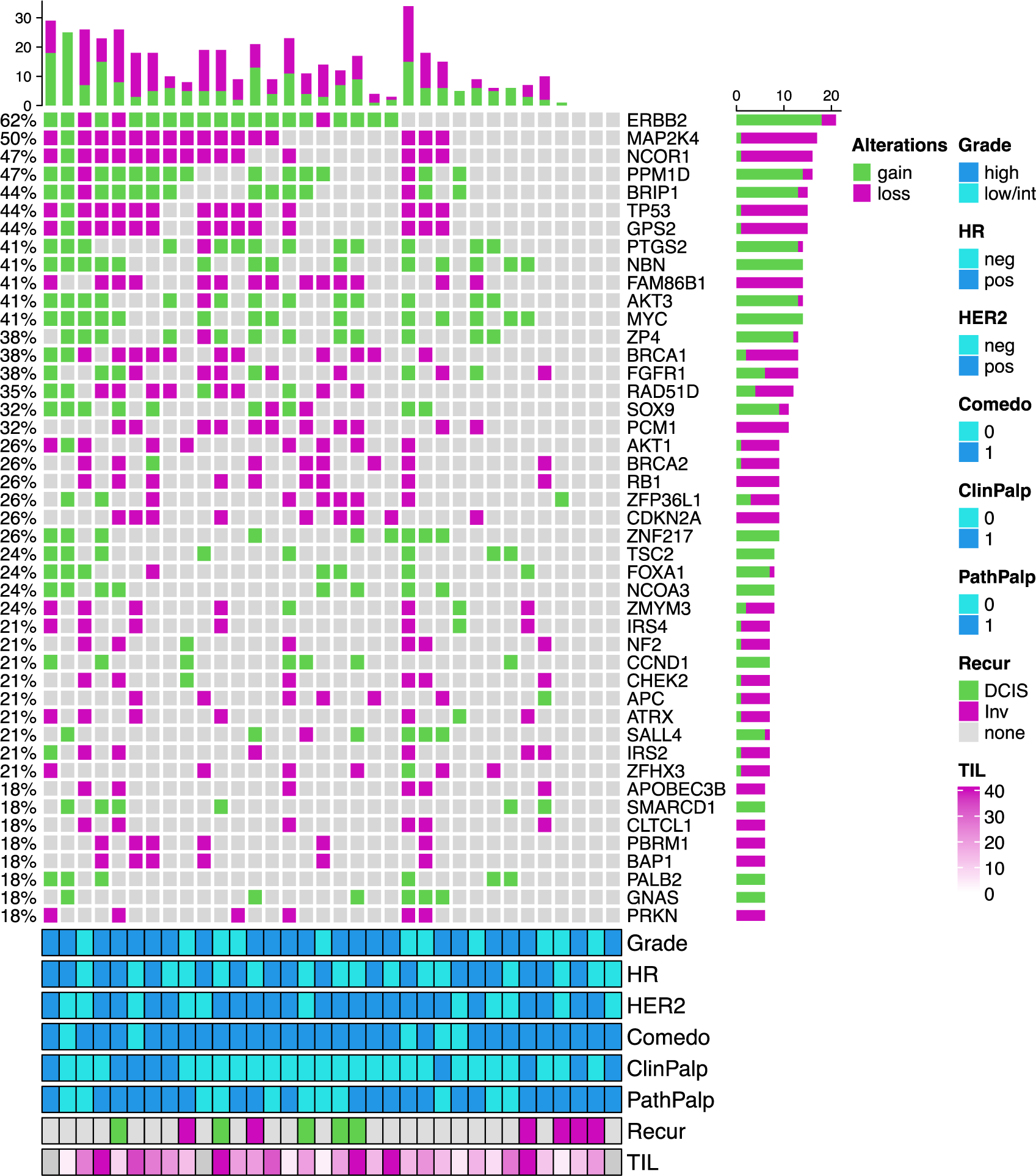
Oncoprint diagram displaying copy number status of driver genes commonly altered in breast cancer. Each column represents a case of DCIS. Clinical characteristics and percent TILs are indicated in the bottom annotations.

**Supplementary Figure S5.**
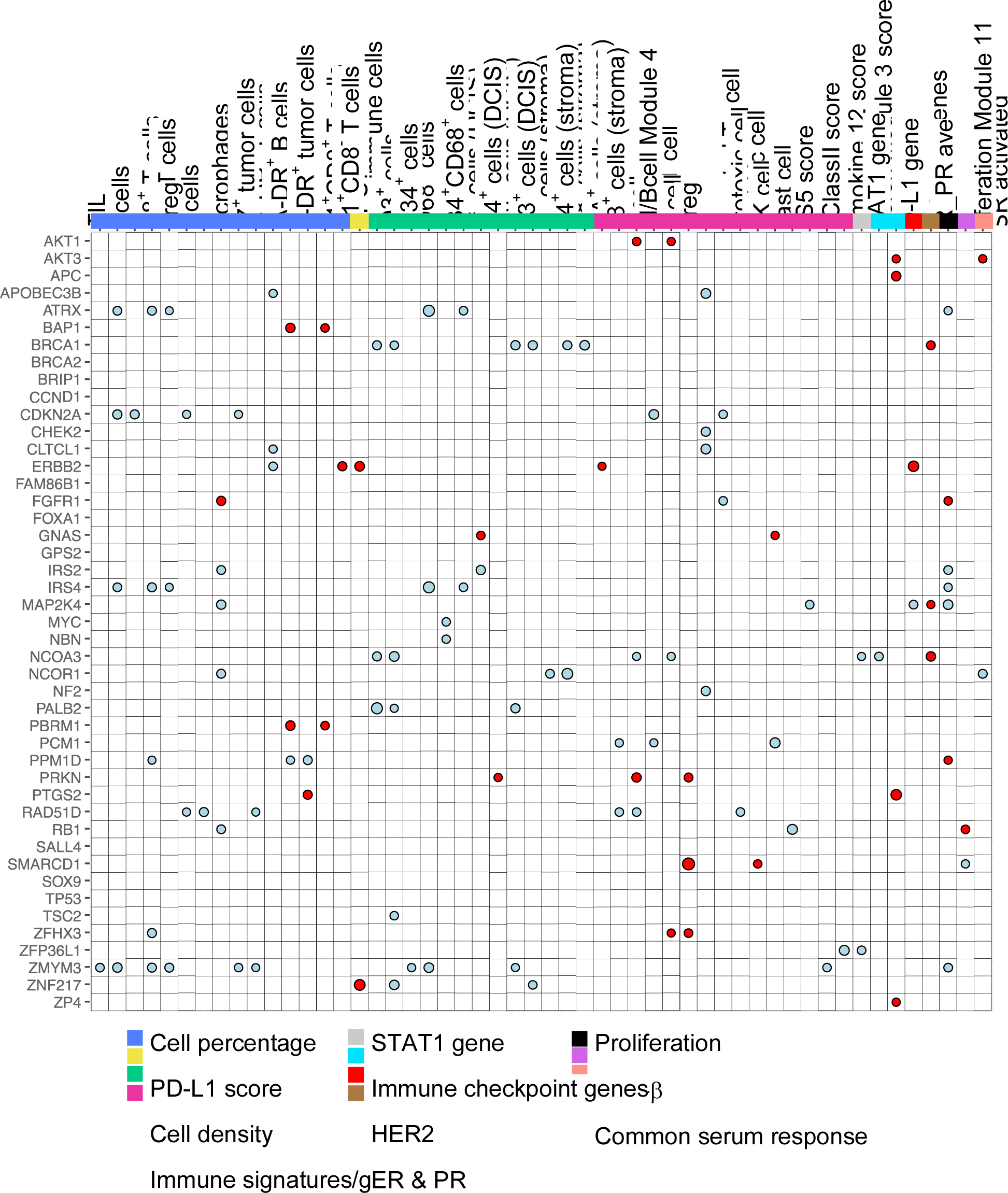
Associations of driver gene copy number alterations with immune biomarkers. Color of dot indicates association with gains (red) or losses (blue). Size of dot is proportional to significance (larger dot → smaller p-value). Only those biomarkers that were significant (p<0.05) with at least one chromosomal aberration are shown. Dots are only shown for significant associations (p<0.05; Kruskal-Wallis test).

**Supplementary Figure S6.**
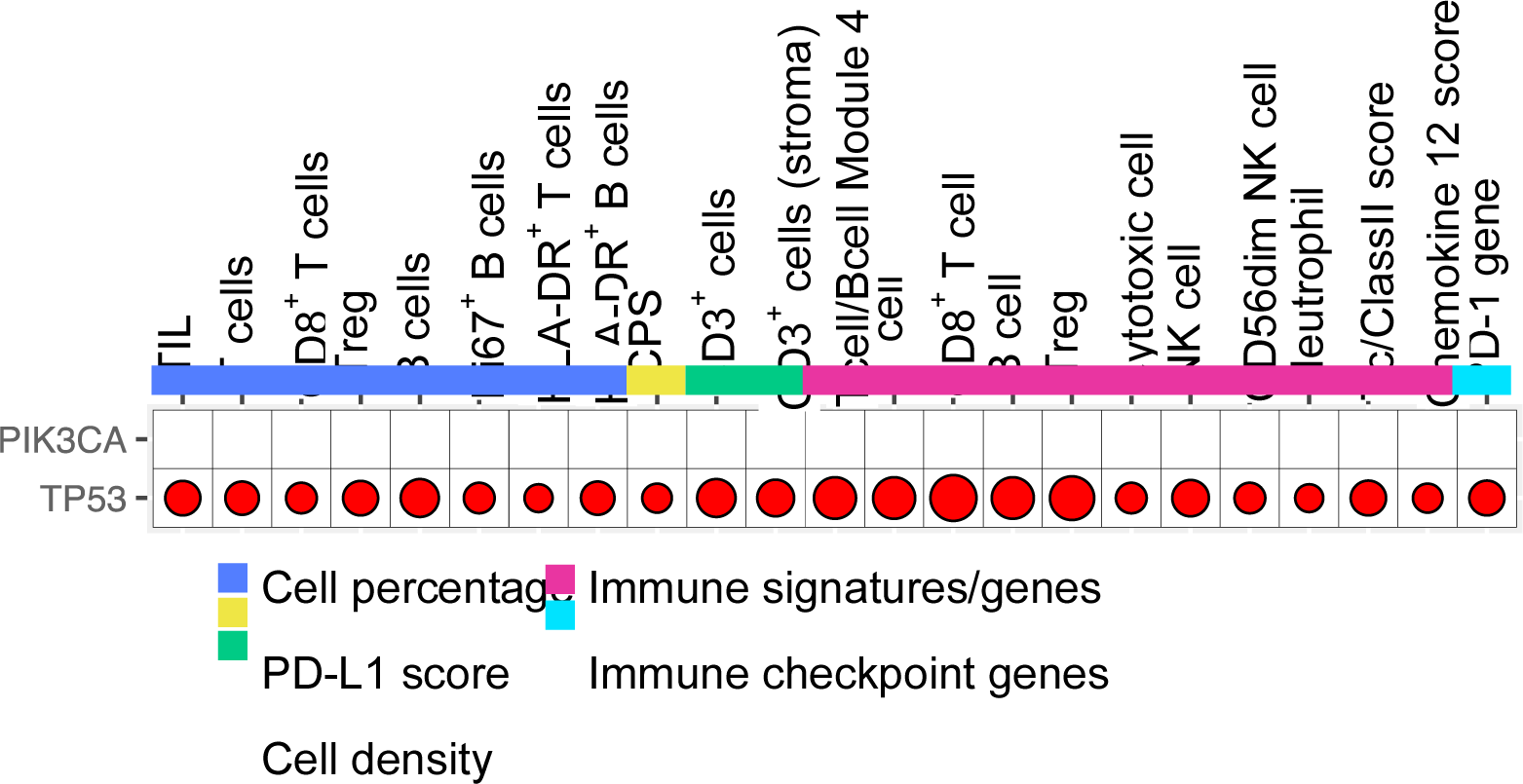
Associations of PIK3CA and TP53 mutations with immune biomarkers. Color of dot indicates direction of association (red: positive, blue: negative). Size of dot is proportional to significance (larger dot → smaller p-value). Dots are only shown for significant associations (p<0.05; Wilcoxon test).

**Supplementary Figure S7.**
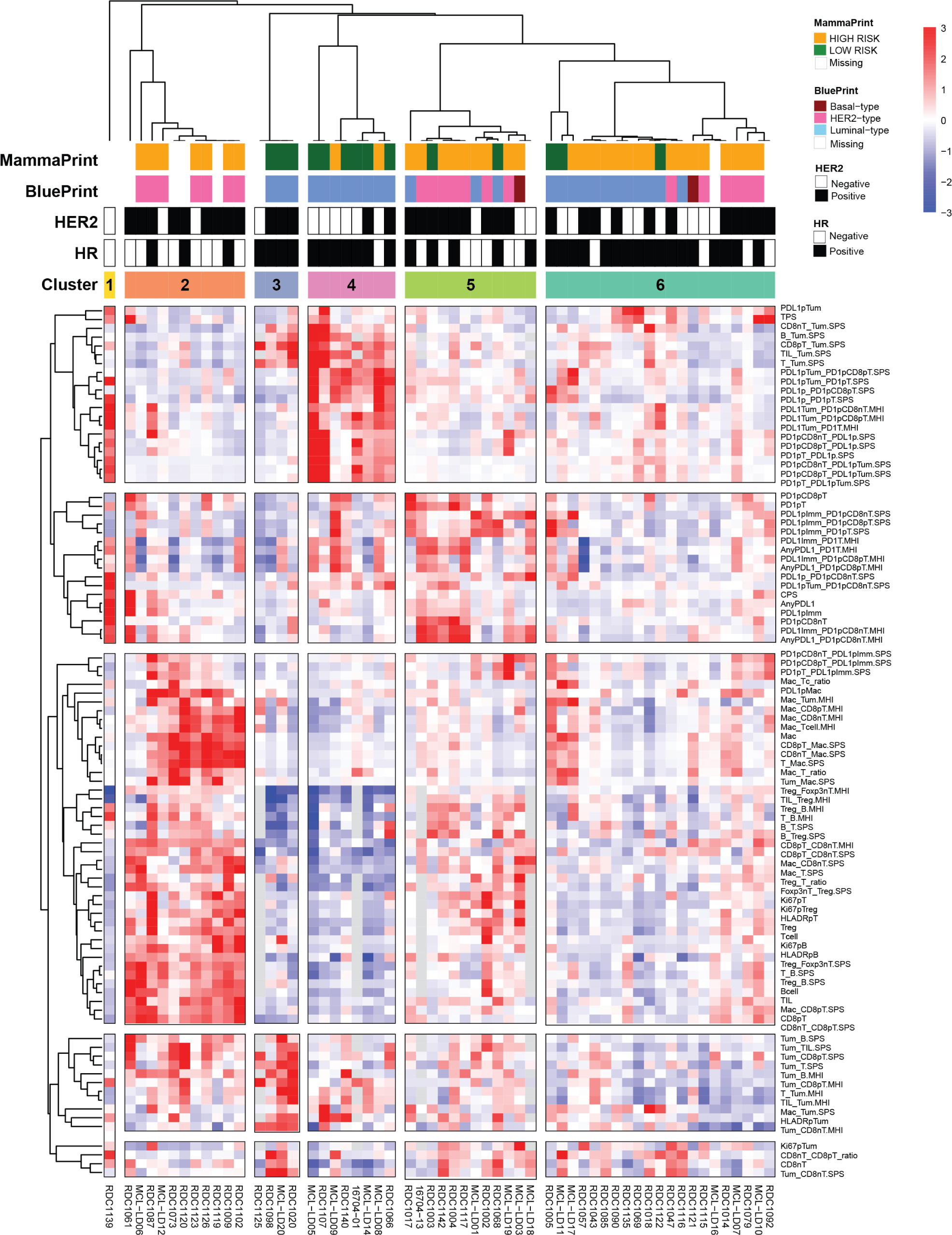
Unsupervised clustering of immune biomarker profiles. Heatmap of median-centered, scaled levels (red: positive, blue: negative) of 94 immune biomarkers assayed on IP1.1 and IP1.2 across 57 samples. Samples are organized along the columns and biomarkers along the rows. Samples are ordered based on consensus clustering (k=6); biomarkers are organized by hierarchical clustering (of median-centered levels) with correlation as distance and average linkage (k=5). Column annotation indicates the MammaPrint Result (Low Risk: Dark green, High Risk: Orange), the BluePrint Subtyping Result (Basal: Dark red ; HER2: hotpink ; Luminal: cornflowerblue), HR/HER2 status (Positive: black, Negative: white) and consensus cluster membership.

**Supplemental Table S4.**
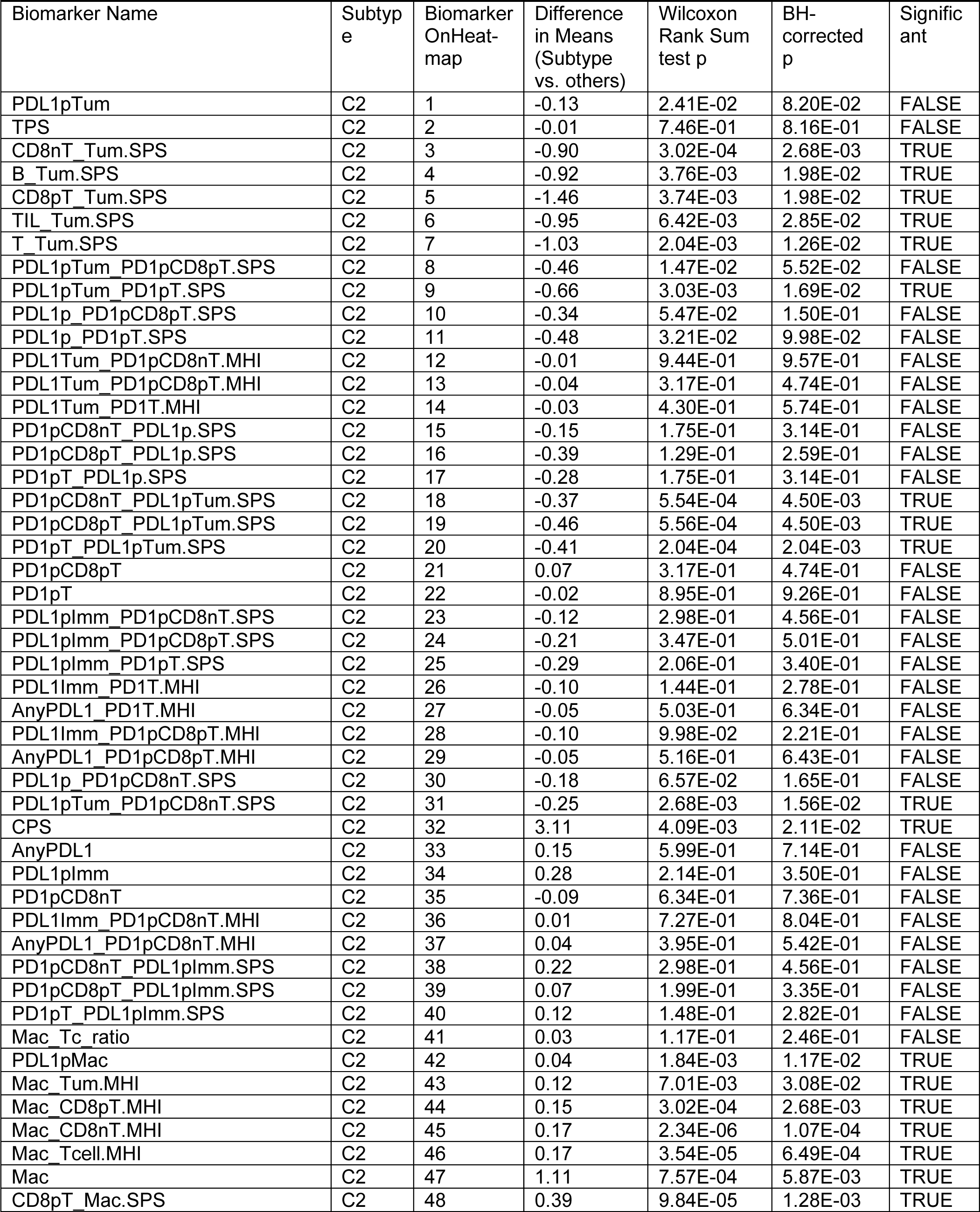
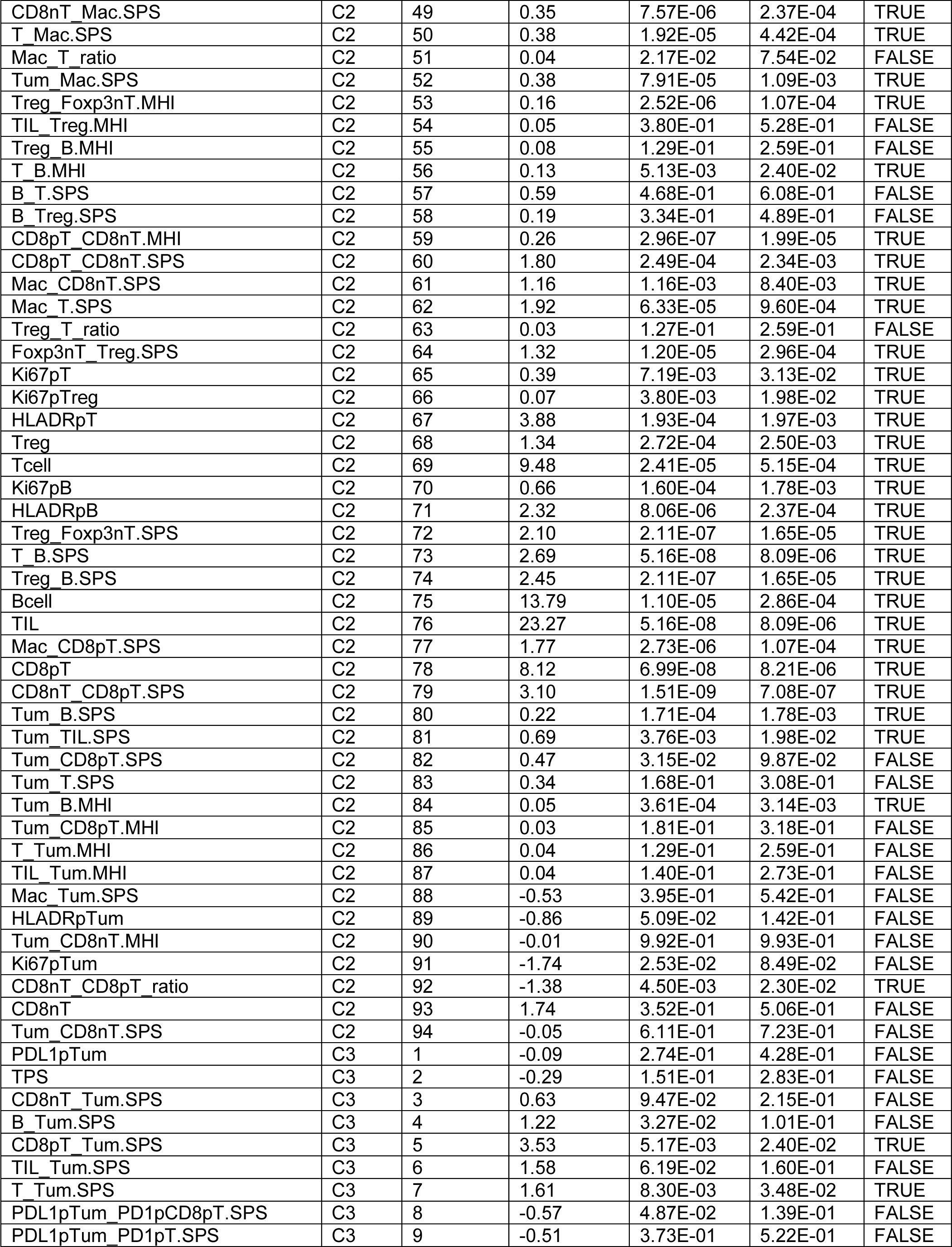
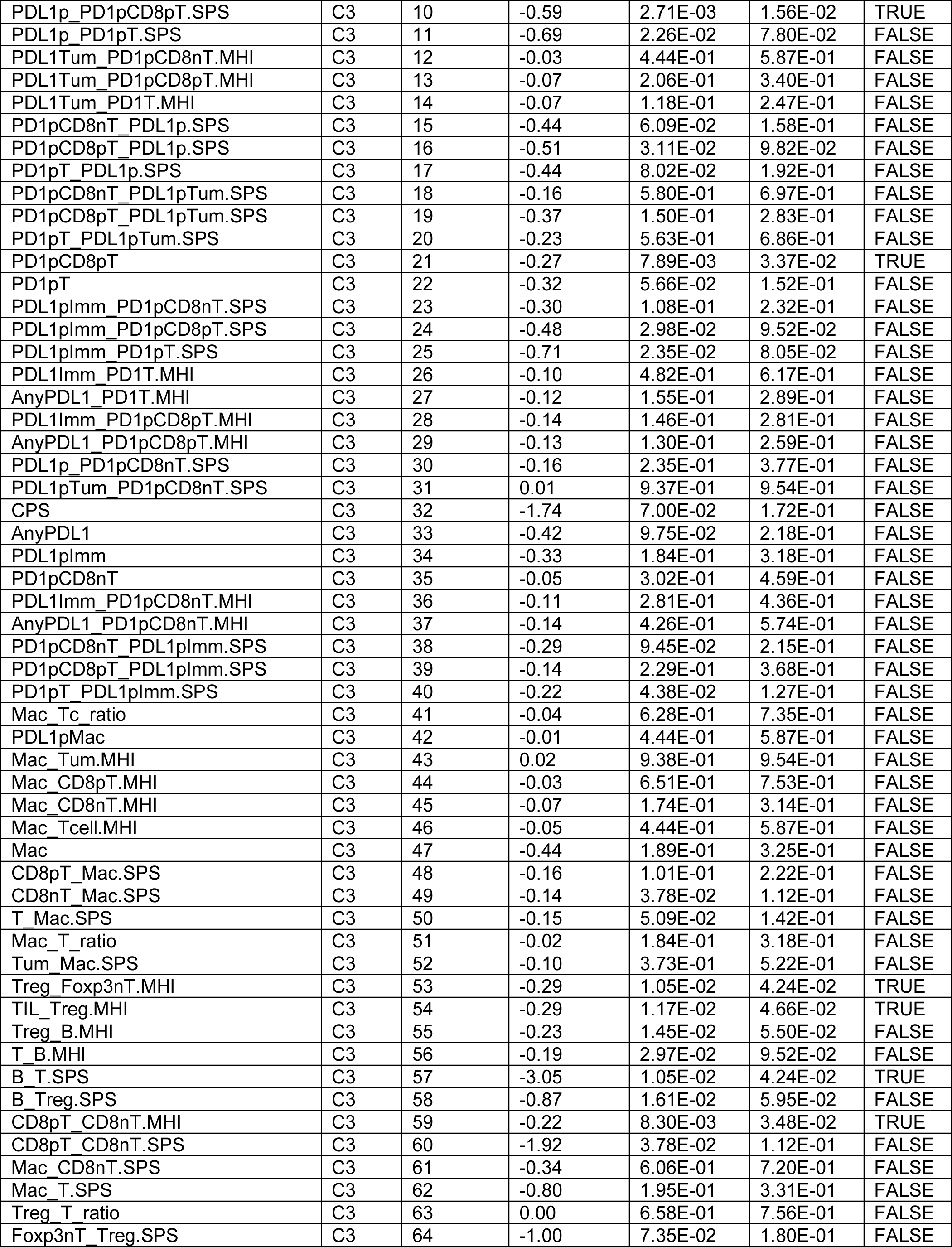
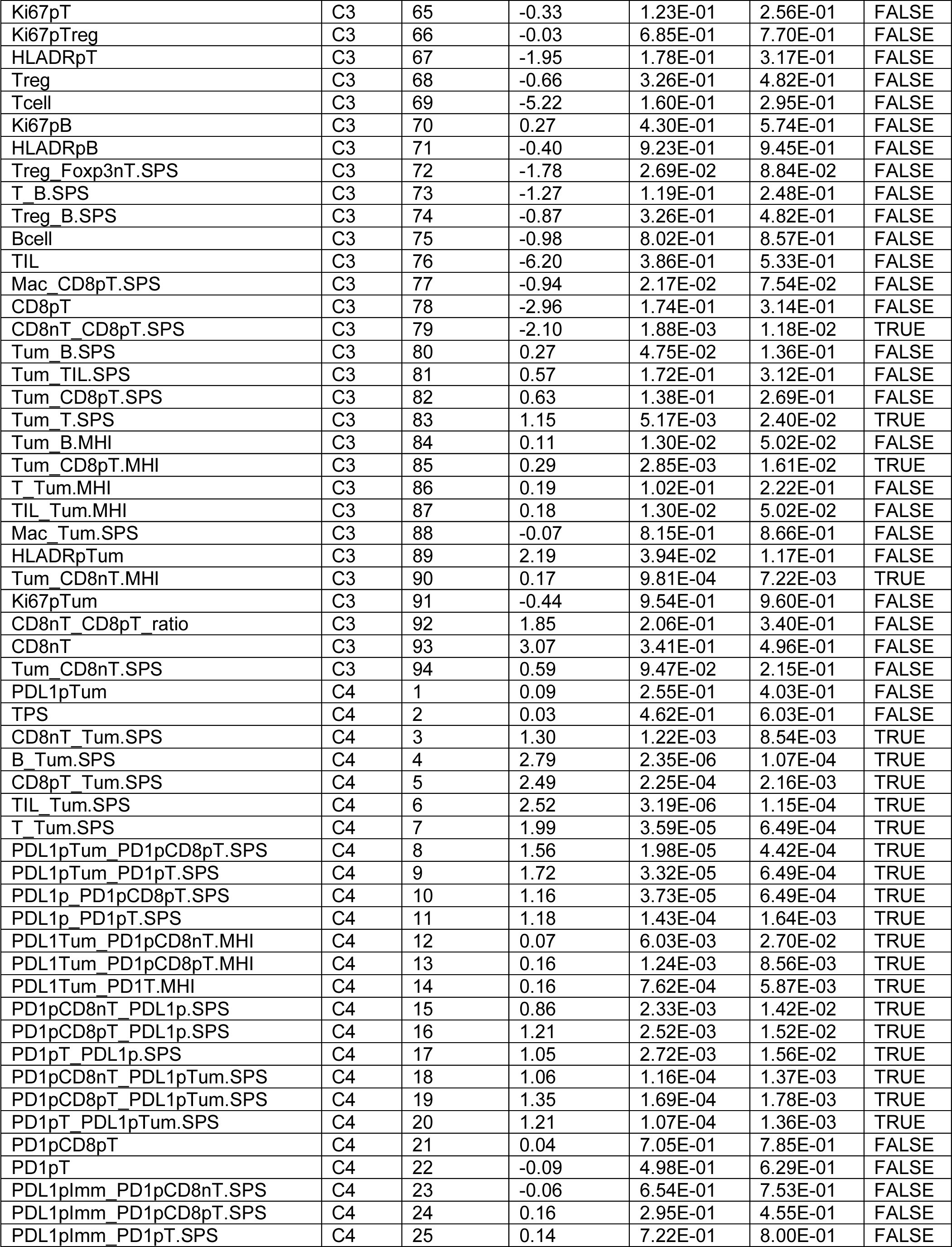
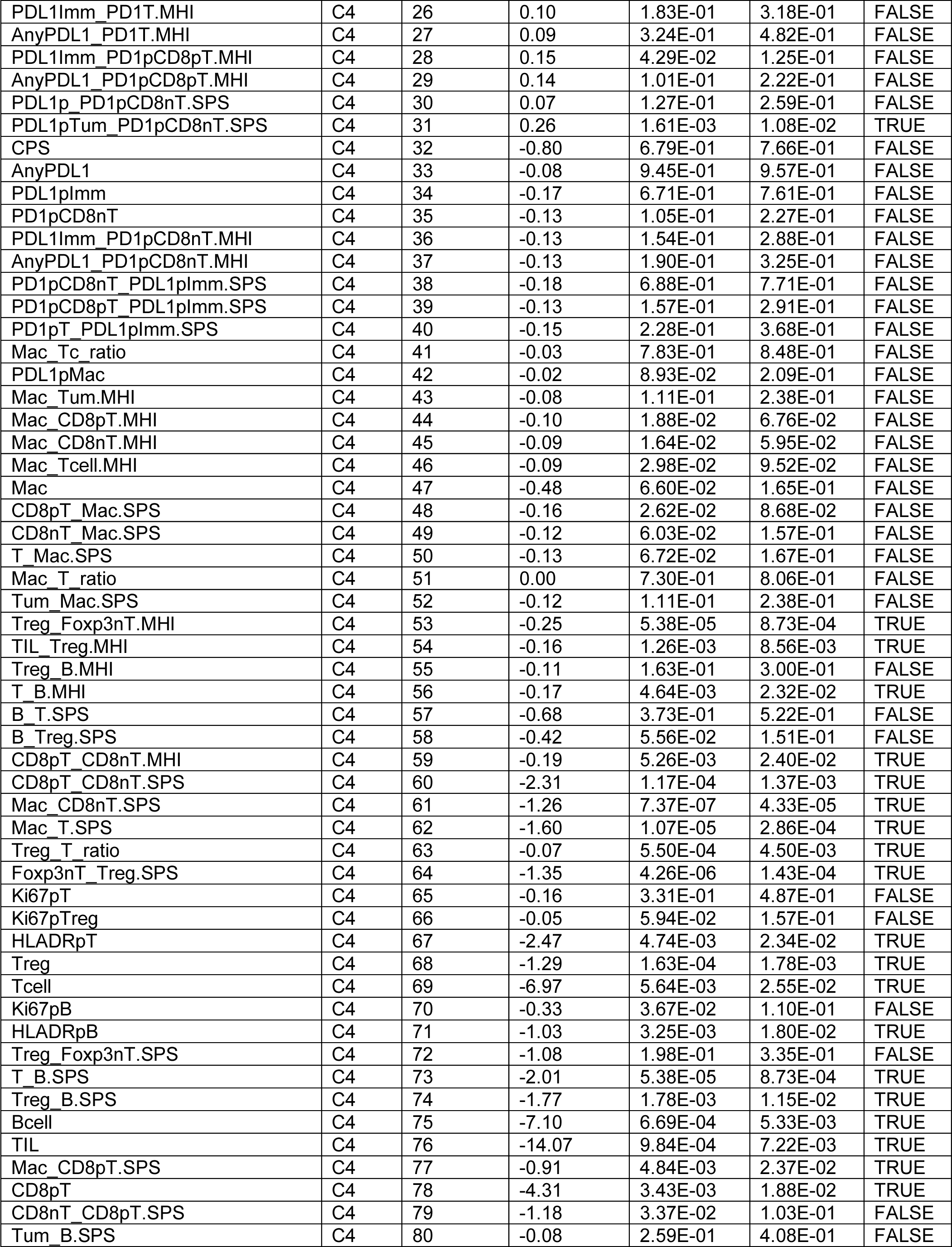
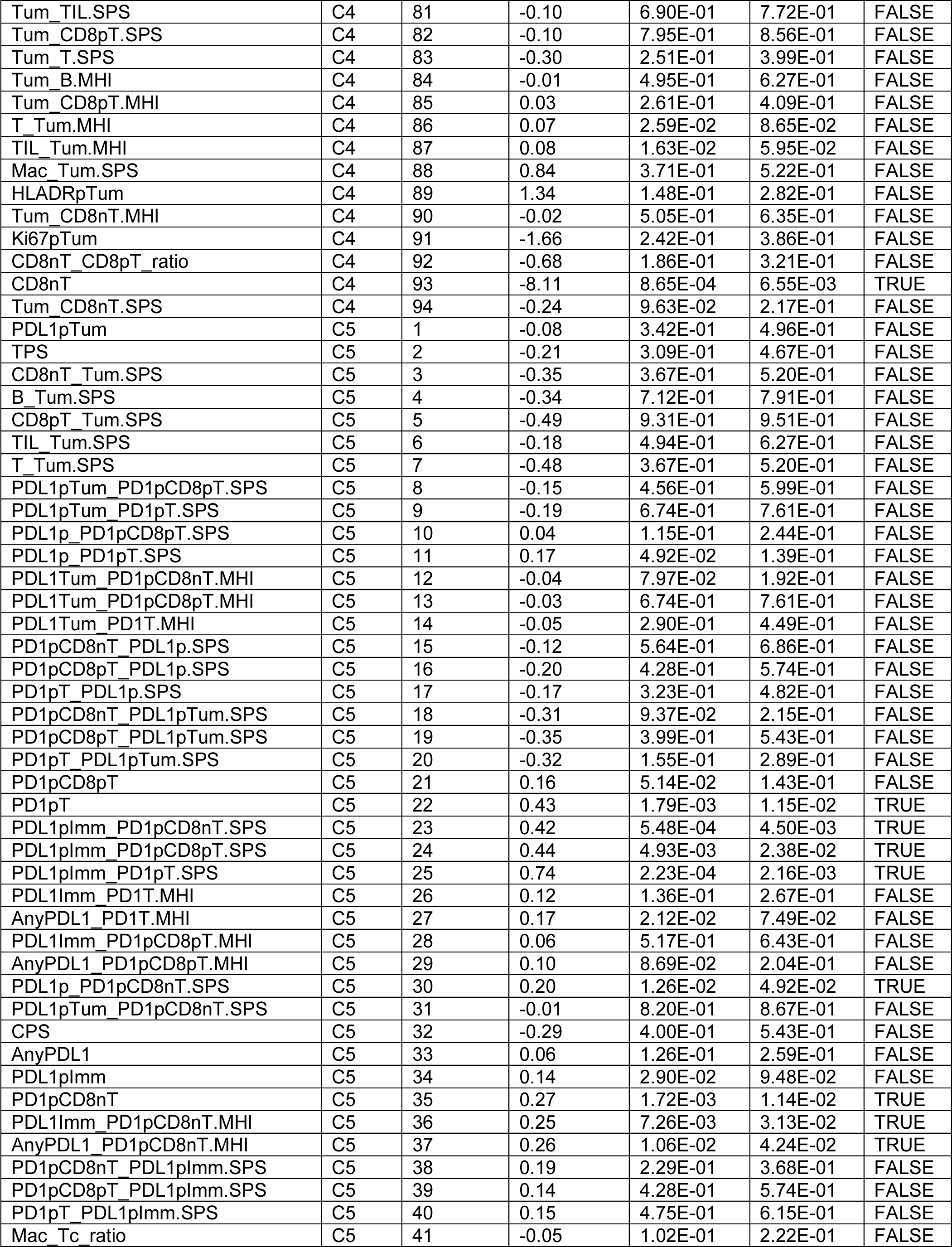
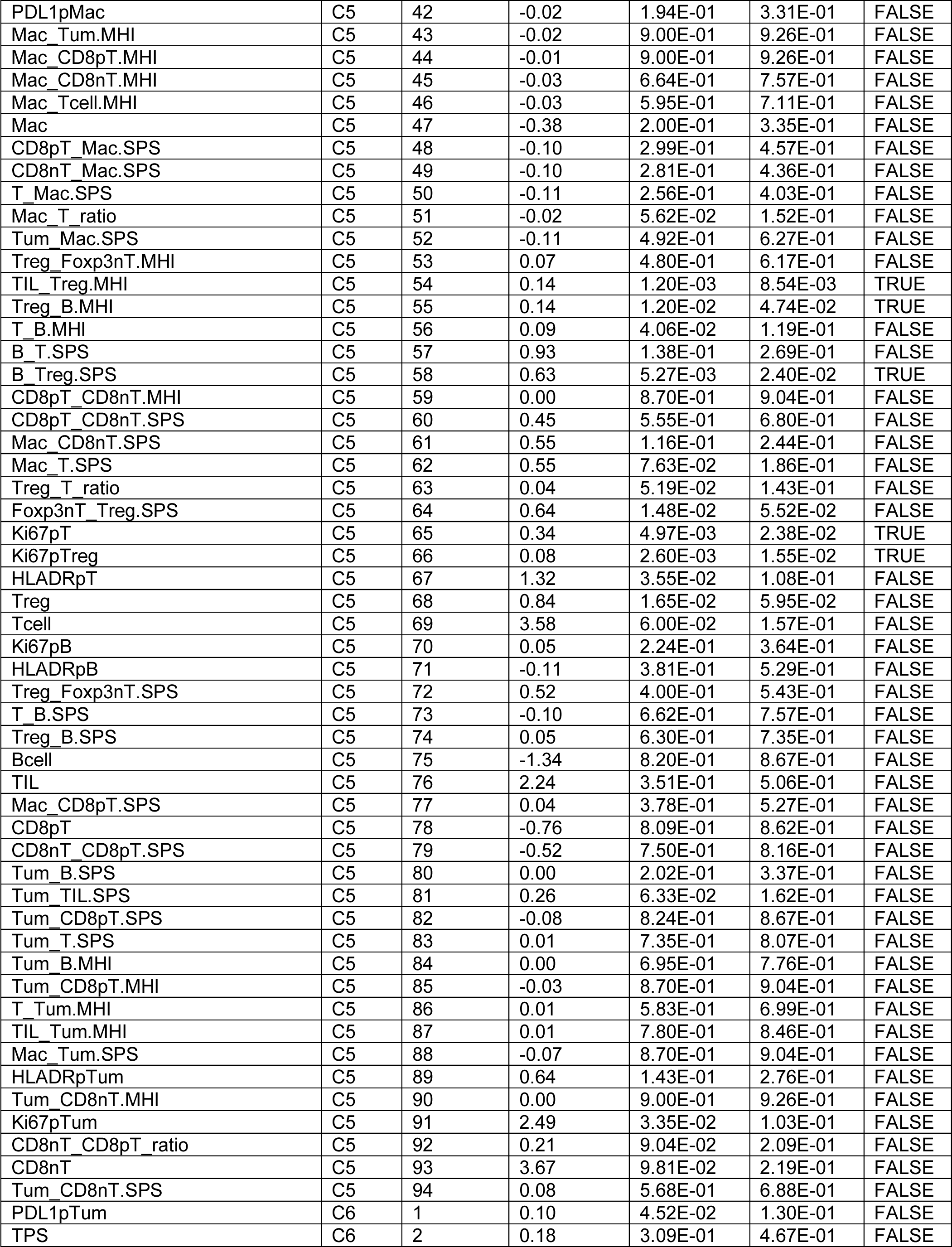
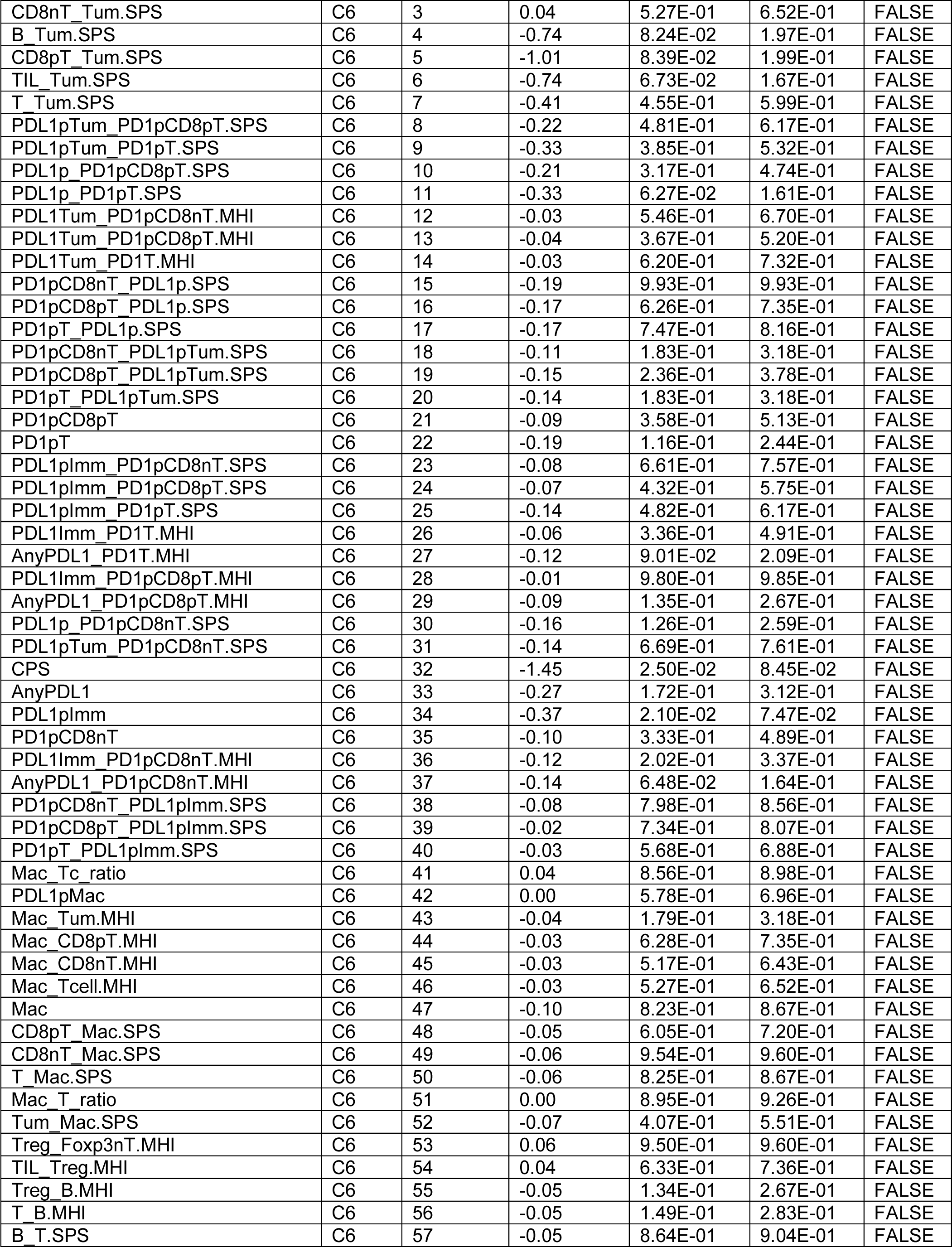
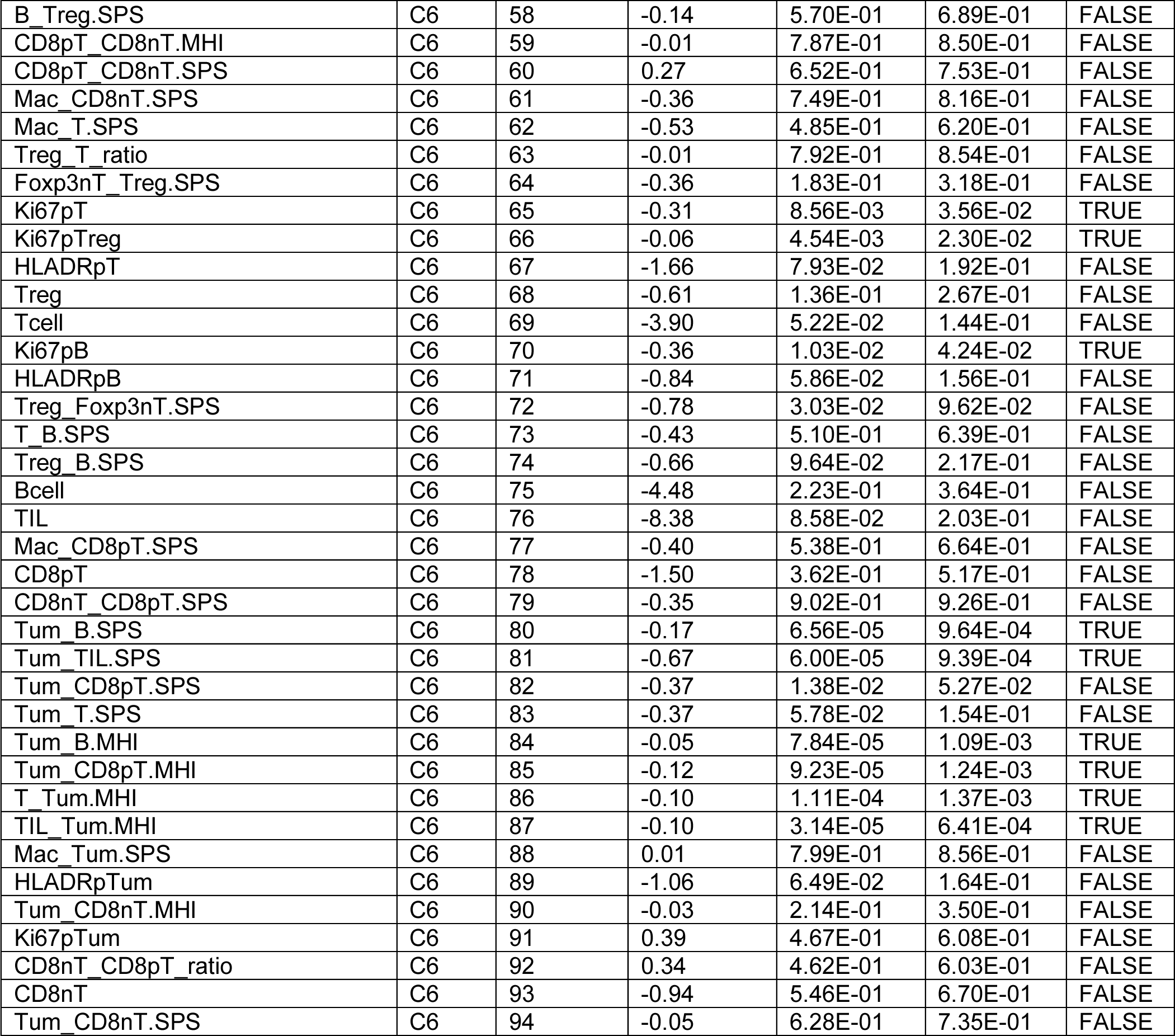
Comparison of immune biomarkers generated on IP1.1 and IP1.2 between DCIS subtypes.

**Supplementary Figure S8.**
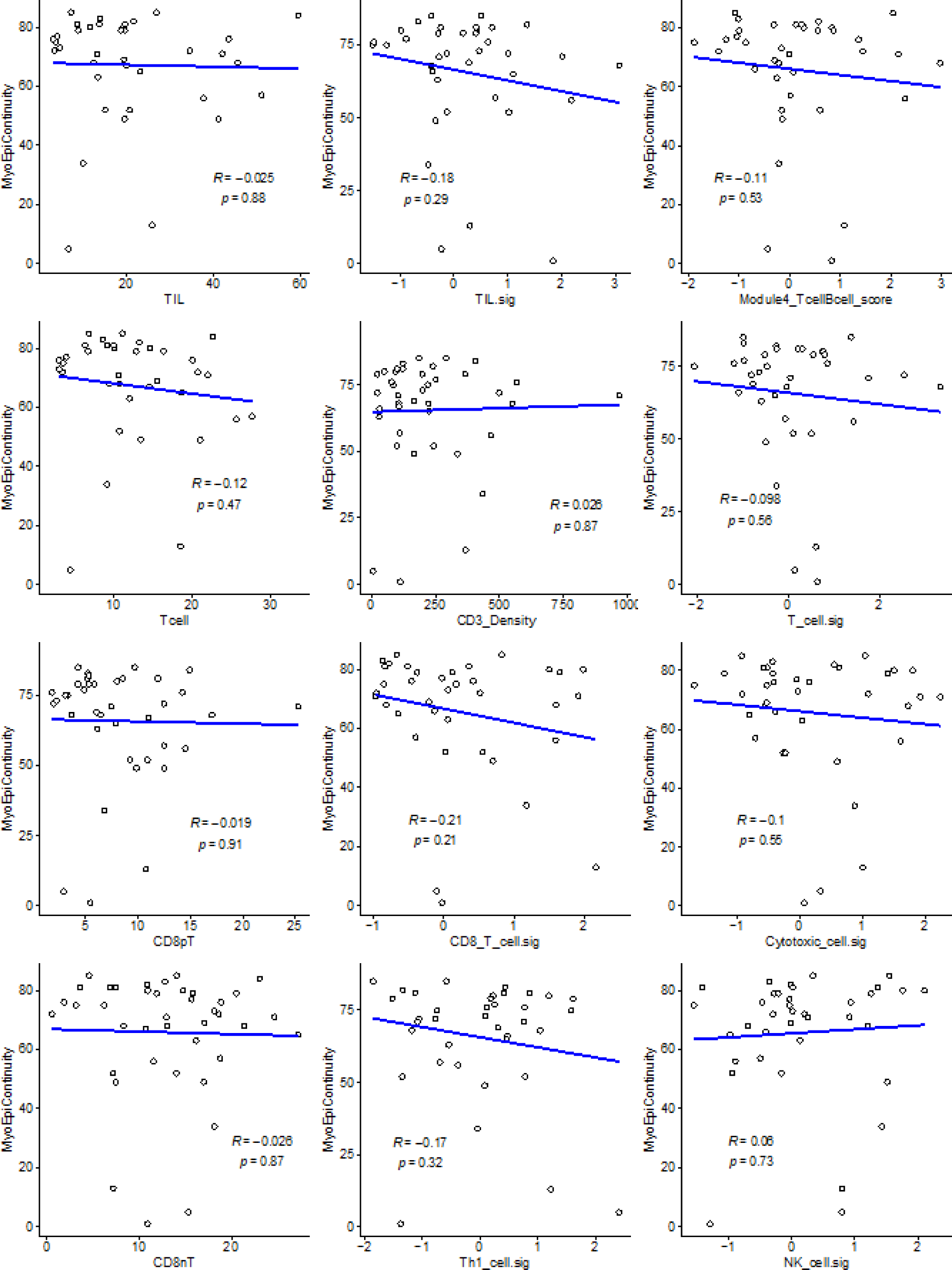
Correlations between cell populations identified by mIF staining or gene expression signatures (x-axes) and myoepithelial continuity score (y-axes). Pearson R and p values

**Supplementary Figure S9.**
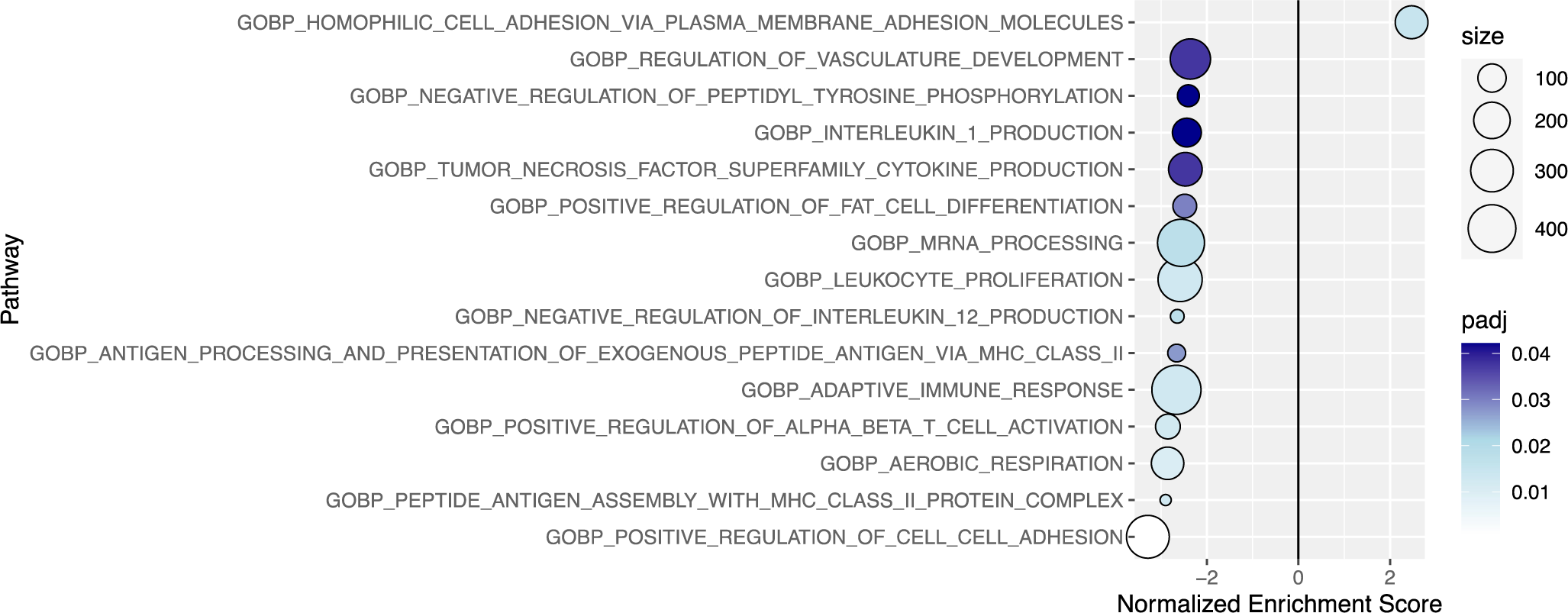
Gene set enrichment analysis (GSEA) identifies ontologies enriched in DCIS with low or high myoepithelial continuity. Normalized enrichment scores are given for each feature ontology. Only ontologies with adjusted p values <0.05 are shown. Size of points is related to the number of genes in each pathway. Points are colored by significance (adjusted p values).

